# Standardized microgel beads as elastic cell mechanical probes

**DOI:** 10.1101/290569

**Authors:** S. Girardo, N. Träber, K. Wagner, G. Cojoc, C. Herold, R. Goswami, R. Schlüßler, S. Abuhattum, A. Taubenberger, F. Reichel, D. Mokbel, M. Herbig, M. Schürmann, P. Müller, T. Heida, A. Jacobi, J. Thiele, C. Werner, J. Guck

## Abstract

Cell mechanical measurements are gaining increasing interest in biological and biomedical studies. However, there are no standardized calibration particles available that permit the cross-comparison of different measurement techniques operating at different stresses and time-scales. Here we present the rational design, production, and comprehensive characterization of poly-acylamide (PAAm) microgel beads mimicking biological cells. We produced mono-disperse beads at rates of 20 – 60 kHz by means of a microfluidic droplet generator, where the pre-gel composition was adjusted to tune the beads’ elasticity in the range of cell and tissue relevant mechanical properties. We verified bead homogeneity by optical diffraction tomography and Brillouin microscopy. Consistent elastic behavior of microgel beads at different shear rates was confirmed by AFM-enabled nanoindentation and real-time deformability cytometry (RT-DC). The remaining inherent variability in elastic modulus was rationalized using polymer theory and effectively reduced by sorting based on forward-scattering using conventional flow cytometry. Our results show that PAAm microgel beads can be standardized as mechanical probes, to serve not only for validation and calibration of cell mechanical measurements, but also as cell-scale stress sensors.

**Significance Statement:** Often vastly different cell mechanical properties are reported even for the same cell type when employing different measurement techniques. This discrepancy shows the urgent need for standardized calibration particles to cross-compare and validate techniques. Microgel beads can serve this purpose, but they have to fulfil specific requirements such as homogeneity, sizes and elasticities in the range of the cells, and they have to provide comparable results independent of the method applied. Here we demonstrate the standardized production of polyacrylamide microgel beads with all the features an elastic cell-mimic should have. These can not only be used as method calibration particles, but can also serve as cell-scale sensors to quantify normal and shear stresses exerted by other cells and inside tissues, enabling many new applications.

## INTRODUCTION

It has become increasingly obvious that cell mechanical properties can be used to monitor physiological and pathological changes in cells, without resorting to molecular markers(1–4). Thus, various technologies have been developed to measure mechanical properties. These are based on the principle that the application of a defined force (stress) can induce a measurable deformation (strain) in cells(5, 6). Such measurements can be carried out using atomic force microscopy (AFM)(7), micropipette aspiration(8), magnetic twisting cytometry(9), particle-tracking microrheology(10), microplate manipulation(11), cell monolayer rheology(12), and optical stretching(13, 14) with characteristic measurement times ranging from tens of milliseconds to minutes. AFM based-indentation is the most commonly used technique to determine the mechanical properties of adherent cells(6, 15–18). The Young’s modulus is typically the mechanical parameter extracted and values reported span a wide range even for the same cell type. This has been recently addressed comparing measurements done on the same cell line (MCF-7) in different labs by using six different techniques(19). The discrepancy of several orders of magnitude was largely attributed to the difference in stress application, timescale, and adherent vs. suspended state. The situation has become even more difficult by the recent advent of high-throughput microfluidic techniques with measurement timescales of milliseconds and below, such as deformability cytometry (DC)(20), suspended micro-channel resonator(21), micro-constriction array(22), real-time DC (RT-DC)(23), and quantitative DC(24). Among them, RT-DC is currently one of the technologies able to induce rapid sample deformation on a time scale of milliseconds and to extract the Young’s modulus(25, 26). Deformability cytometry(20) can even reach timescales of microseconds, but, due to its operation in the high Reynolds number regime, calculations of cell elasticity are generally hard and have not been demonstrated yet.

Within this large number of available techniques to measure cell deformability, AFM indentation and RT-DC serve well to point out the differences among them. AFM indentation and RT-DC operate at different timescales (10^−2^ – 10^1^ s vs. 10^−3^ s) and force levels (0.5 – 10 nN vs 100 nN – 1 µN), respectively, which both can lead to different elasticity values due to the cells’ time-dependent and nonlinear mechanical properties(25, 27). Furthermore, measurements on cells by AFM and RT-DC are done in different conditions, respectively on adherent and non-adherent cells. Thus, it is hard to identify the cause(s) of the observed differences in elasticity measured by using these two techniques. Importantly, these aspects cannot be investigated using cells as probes due to their complexity, biological variation, and non-linear viscoelastic properties(19, 28). Therefore, homogeneous synthetic particles with sizes and elasticities in the range of the cells being investigated are needed to compare different techniques. To reach comparable results independent of the method applied, it is also necessary that such particles have timescale-independent mechanical properties (i.e., a purely elastic material).

Agarose particles have been used to calibrate novel microfluidic techniques, such as quantitative DC and RT-DC. The beads were produced by using emulsification. Mietke *et al.* measured by RT-DC a Young’s modulus of 2.2 ± 1.1 kPa for agarose beads with an agarose concentration of 0.5% (w/w)(25). A similar Young’s modulus, 2.0 ± 1.7 kPa, was measured by Nyberg *et al.* using AFM, but for a higher agarose concentration of 2.5% (w/w)(24). Furthermore, the agarose beads showed a high coefficient of variation (50 – 80%) in their elasticity for a fixed composition, suggesting that not all the beads polymerized in the same way. This variability demonstrates that not only cells, but also synthetic particles that should have the same properties, exhibit different Young’s moduli. The measured difference is likely due to viscoelastic energy dissipation in the non-covalently cross-linked agarose gel that had not been considered(29). This influence can be minimized by choosing a different, chemically cross-linked material.

Among the existing options, polyacrylamide (PAAm) has been intensively studied(30, 31) and accepted as an elastic material(32, 33). Nevertheless, various factors can affect its gelation kinetics and network formation (i.e., total monomer concentration, cross-linking monomer percentage, temperature, viscosity)(31, 33, 34). As a result, the final gels are far from the “ideal” gel concept of Richards and Temple(30), and different defects can be included in the polymer network randomly. This affects their physical properties such as swelling, elasticity, transparency, and permeability, leading to different elastic moduli despite starting from the same polyacrylamide composition(31). These aspects have been extensively studied on bulk gels, but have never been analyzed, or attempted to be controlled, on microgel beads with a size comparable to cells.

Here we demonstrate the reliable production of PAAm microgel beads as standardized calibration samples for cell mechanical measurements. Droplet microfluidic technology was used for the high-throughput (20 – 60 kHz) production of microgel beads with controlled size and elasticity, keeping the cross-linker to monomers ratio concentration constant and changing the total monomer concentration. Optical diffraction tomography and confocal Brillouin microscopy demonstrated that beads were homogeneous at the microscale. This important verification of a central assumption of the Hertz model enabled a rigorous and reliable analysis of their elasticity by AFM, demonstrating the ability to cover an elasticity range of interest for cell mechanics. Based on polymer physics, the elasticity values and swelling behavior were used to calculate structural parameters of the gel, such as cross-link density and porosity. Knowledge of these parameters allowed us to understand the origin of elasticity variation and to investigate the poroelastic response of the gel. The beads were further analyzed by RT-DC, which operates at much shorter timescales but yielded comparable elastic moduli. As an intriguing option, we demonstrated that the remaining variability in elasticity can be significantly reduced by sorting the beads based on their forward light scattering properties using a conventional flow cytometer. Finally, beads were functionalized with Poly-L-lysine (PLL) for non-specific cell attachment and fluorescence, and embedded in multicellular aggregates. Their interaction with the surrounding cells was demonstrated by their deformation during aggregate growth. These fully characterized beads do not only represent the first example of a standardized sample for validation and calibration of cell mechanical measurements. They also open new perspectives towards their use as passive cell-scale sensors able to interact with biological samples and to sense acting stresses through their deformation.

Furthermore, our study provides a combination of high-throughput technologies that can be used for the fast production, characterization, and sorting of novel smart microgel probes.

## RESULTS

### PAAm microgel bead production and homogeneity

Microgel beads with a diameter of <20 µm were produced to match the typical size of eukaryotic cells being analyzed by AFM indentation and RT-DC. In general, there are various technologies available that can control shape and size of microgels, such as micromolding, photolithography, extrusion, emulsification and droplet microfluidics(35). Among them, droplet microfluidics is able to provide monodisperse droplet microreactors with high throughput(36, 37). A flow-focusing PDMS-based microfluidic device was used to generate the microgel particles(38), where the flow of the dispersed and continuous phase was controlled by a pressurized pump. In the device, the main stream of the dispersed phase, containing the pre-gel mixture, was squeezed by the two lateral flows of the continuous oil phase (inset of Figure 1A). This led to the formation of pre-microgel droplets, which were then polymerized by the free-radical crosslinking copolymerization of acrylamide (AAm) and N,N′-methylenebisacrylamide (BIS) initiated by ammonium persulphate (APS). The volume of the pre-gel mixture, *V_sol_*, containing AAm, BIS and APS was fixed at 100 µl, and the time necessary to consume it, *t_c_*, was recorded. Different pre-gel solutions were prepared by varying the total monomer concentration defined as between 5.9 – 11.8%, but maintaining the cross-linker to monomer ratio concentration constant at *C* = 3.25%.

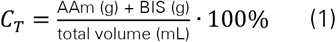

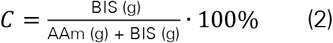

**Figure 1.**
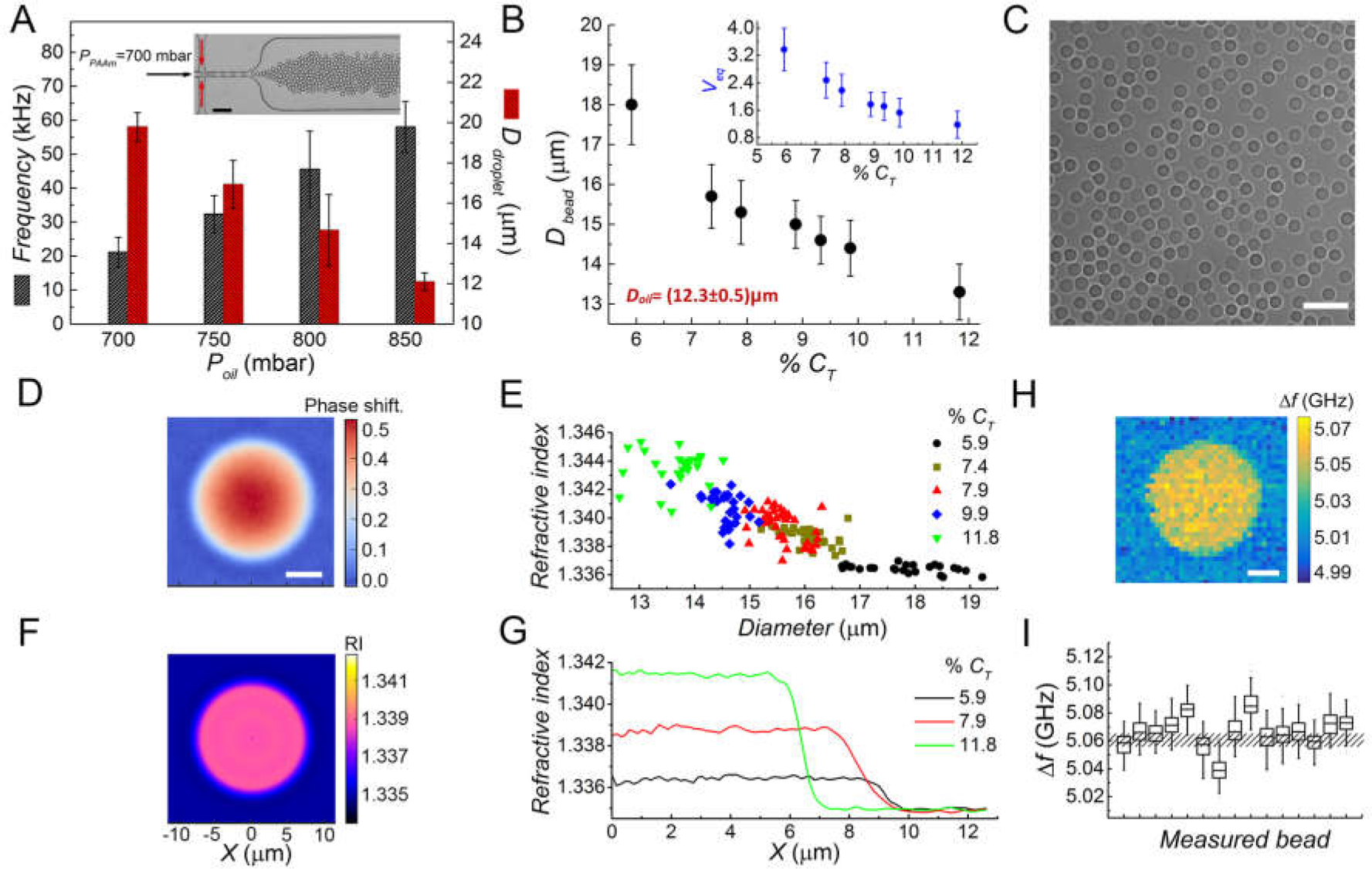
Production, swelling and homogeneity of PAAm beads. (A) Influence of oil pressure *P_oil_* change on PAAm droplet size and production rate during fabrication (*n* = 3) for *C_T_* = 9.9%. Inset: bright-field image of flow focusing microfluidic device during PAAm droplet production. Movie available in S.I. Oil with surfactant and TEMED (red arrows) is injected perpendicular to the polyacrylamide pre-gel solution (black arrow) and result in the formation of PAAm droplets at the junction. Data are shown for constant polymer pressure of 700 mbar and *C_T_* = 9.9%. Scale bar: 50 µm. (B) Final PAAm beads diameter in 1x PBS in dependence on *C_T_*, after overnight swelling. The swelling behavior of the beads is shown for an initial PAAm droplet diameter of (12.3 ± 0.5) µm, obtained with 850 mbar oil pressure. The inset shows the swelling behavior of the beads displayed as the normalized volume of equilibrium of the swollen gel *V_eq_* in dependence on *C_T_*. Data is also shown in Table S1. (C) Bright-field image of polymerized PAAm beads in 1x PBS. Scale bar: 50 µm. (D) Quantitative phase image of a PAAm bead (*C_T_* = 7.9%) for refractive index (RI) determination. Color scale bar: Phase shift in radians. Scale bar: 5 µm. (E) RI variation with polymer composition and bead size. Data points indicate the RI for *C_T_* ranging from 5.9% – 11.8%. Each point corresponds to a single measured bead. (F) Cross section of a tomographic RI reconstruction from a single quantitative phase image (by assuming spherical symmetry) of a PAAm bead. Color bar: RI values. (G) Homogeneity of RI distribution for 11.8% (green line), 7.9% (red line) and 5.9% (black line) *C_T_* obtained from radial RI profiles. (H) Brillouin shift (*Δf*) image of a single bead (*C_T_* = 7.9%). The variation of Brillouin shift within the bead corresponds to the random error of the measurement system. Scale bar: 5 µm. (I) Brillouin shift of 15 different beads measured with the Brillouin microscope (*C_T_* = 7.9%). The width of the patterned area shows the intensity of the systematic error of the system (10 MHz).

The continuous oil phase was a solution of fluorinated oil (HFE-7500) containing an ammonium Krytox® surfactant to stabilize the emulsion, and TEMED to catalyze the polymerization process in the droplets. The surfactant was synthetized and further characterized in solution with respect to emulsion stability, infrared (IR) spectrum, and critical micellar concentration (Figure S1). The pre-gel solution pressure was kept constant (700 mbar), while oil pressure was varied between 700 to 850 mbar, which lead to an increase in droplet production frequency from 21,000 to 58,000 droplets/s and a decrease in droplet size from 20 to 12 µm (Figure 1A)(37). The droplet production process (size and frequency) was independent of the total monomer concentration. The diameter of 20,000 droplets was measured by brightfield microscopy and was not varying with the pre-gel mixture composition (Figure S2-A). The mean droplet size, 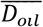, used here was fixed at about 12 µm, and polymerization was carried out in the oil phase at a temperature of 65 °C. Although gelation is expected to occur within 400 min(39), the droplets were left overnight to allow reaction to completion. Finally, the emulsion was broken, and the size of the microgel particles re-suspended in 1x PBS was analyzed, after overnight swelling. All beads were found to be of uniform size, with a coefficient of variation lower than 5.5% (Table S1). The mean diameter of the final beads, 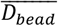, varied from 13.3 ± 0.7 µm to 18.0 ± 0.9 µm as *C_T_* was decreased from 11.8 to 5.9% (Figure 1B), demonstrating that solvent imbibition into the microgel meshwork depends on gel composition. The swelling behavior of the beads was quantified by calculating the normalized volume of equilibrium swollen gel, 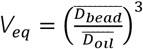, (34) as shown in the inset of Figure 1B by using the data reported in Table S1. The repeatability of the swelling behavior for different batches at three concentrations is shown in Figure S2-B. The final hydrogel beads (Figure 1C) were further characterized by determining their refractive index using quantitative phase microscopy to check if TEMED diffusion from the oil phase to the droplets introduced a radial variation in density(40). Quantitative phase images of the beads were acquired on an inverted microscope equipped with a quantitative phase-imaging camera (Figure1D), and their refractive index (RI) value, which is proportional to mass density, was calculated as described previously(41). The RI increased with increasing *C_T_*, was independent of bead diameter, and reached values between 1.3358 and 1.3454 (Figure 1E). To verify the uniformity of RI distribution within the hydrogel beads, radial RI profiles were computed from representative phase images using optical diffraction tomography(42, 43). A cross-section through the center of the reconstructed volume is displayed in Figure 1F. Figure 1G provides data on RI variation along the bead radius for three different gel compositions, and demonstrates that RI and mass density does not vary along the radial direction. In addition, confocal Brillouin microscopy images through a central plane of the beads also showed no radial variation in Brillouin shift, Δ*f* (Figure 1H). Together, the constant value of the RI and Brillouin shift inside each bead demonstrated that the longitudinal elastic modulus is not varying within the bead volume(44, 45). Thus, the beads produced were isotropic and homogeneous in the investigated range of total monomer concentrations. This result is an important verification of the assumption required to analyze objects by AFM nano-indentation using the Hertz model. Brillouin analysis of different beads, obtained by using the same monomer concentration (*C_T_* = 7.9%), showed a variation in the Δ*f* mean value, suggesting a certain variation in the elasticity of individual beads despite nominally fixed composition (Figure 1I). Taken together, these measurement results were important to understand how at each droplet composition the polymer meshwork is able to take up different amount of water affecting the mechanical and structural properties of the beads(46). A weak polymer network imbibes more water, corresponding to a low value of elastic modulus and refractive index.

### Mechanical characterization by AFM, importance of the analysis model chosen and poroelastic effect

AFM-based indentation (Figure 2A) was used to quantify PAAm microgel bead elasticity. AFM uses a tip of selected geometry to indent an object, and determines the applied force from the bending of the AFM cantilever. Fitting the force-indentation curve to the Hertz model for the corresponding tip geometry can give quantitative measurements of material elasticity, extracting the object’s Young’s modulus, *E*. When the cantilever indents the bead, two effects have to be considered. First, depending on the geometry of the probe and its size compared to that of the object, an additional deformation can be caused by the counter pressure from the substrate at the bottom of the object. This deformation can be neglected for sharp tips(47), but not for a spherical indenter of size comparable to the object. This aspect was analyzed by Dokukin *et al.*(48) and Glaubitz *et al.*(47), introducing a double-contact model for soft spherical samples that extends the conventional Hertz model by a factor *k*, capturing the compression from the substrate, to prevent significant underestimation of the particle’s Youn’s modulus.

**Figure 2.**
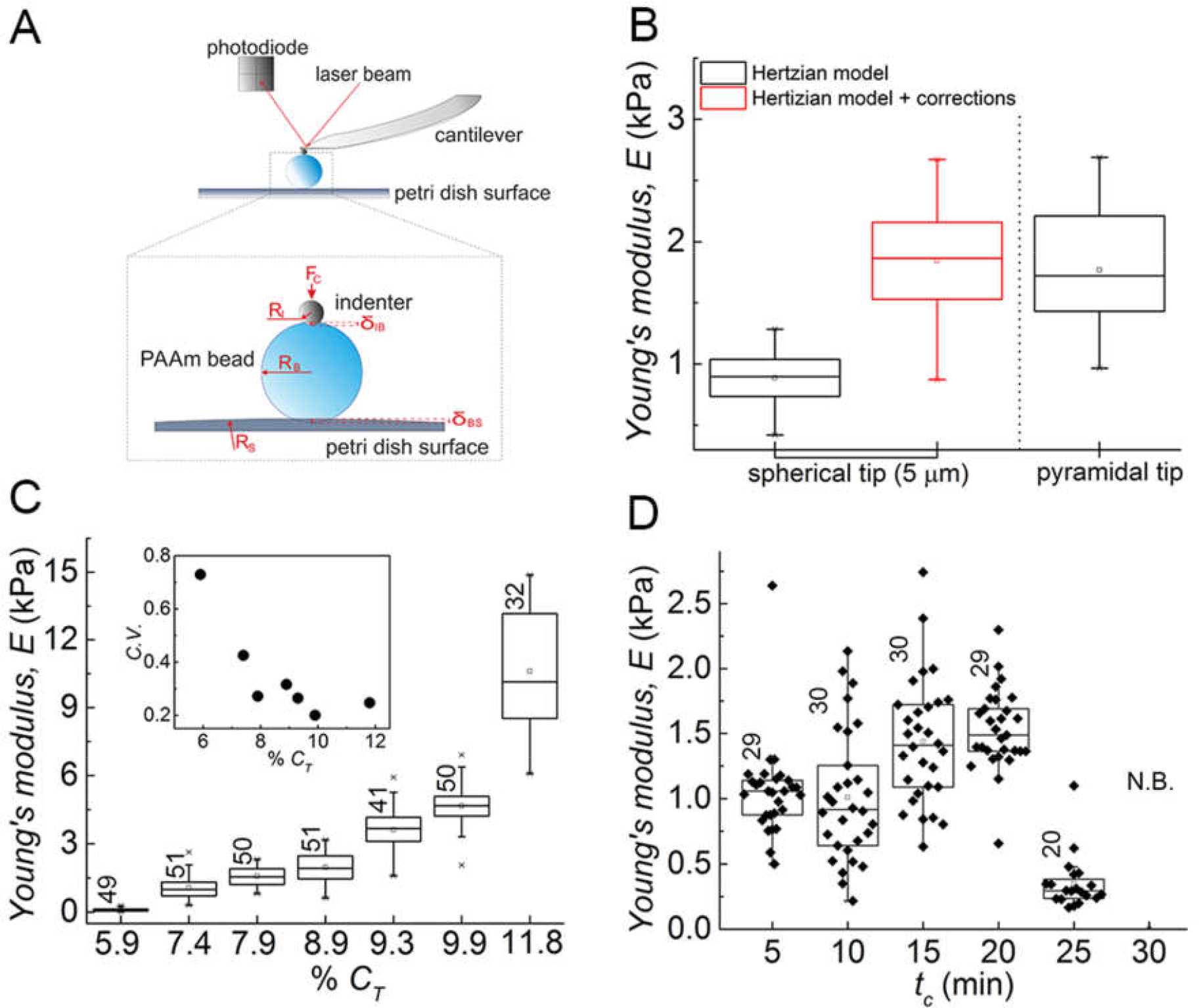
Microgel bead elasticity analysis by AFM indentation. (A) Schematic of the AFM setup. The spherical probe of the cantilever (radius *R_I_*) indents the PAAm bead (radius *R_B_*) with a certain force *FC* while a laser beam is directed on its backside and reflected to a photodiode for the detection of the cantilever deflection. During indentation, the PAAm bead is additionally deformed by the petri dish surface (*RS*) on the bottom region, which requires the application of corrections using the double contact model(47) to determine Youn’s modulus. (B) The depiction of PAAm bead Young’s modulus obtained measuring the same ten beads (*C_T_* = 7.9%) with different indenter geometries (spherical tip: diameter 5 µm, pyramidal tip). Data is shown for the application of the Hertz model with (red box) and without (black boxes) corrections and compared to Youn’s modulus obtained with a pyramidal indenter (n=31). (C) Youn’s modulus variation of PAA beads versus the total monomer concentration. (D) Effect of collection time (*t_C_*) on bead gelation and elasticity during droplet production. No beads (N.B.) were obtained after 25 min.

The second effect to be considered for proper analysis of the force-indentation curves is that in general materials are not just elastic, as assumed in the Hertz model, but that energy can be dissipated. While viscoelastic dissipation can be neglected for PAAm hydrogels in AFM measurements(49), poroelastic effects — the migration of solvent molecules across the gel into the external solvent — have to be considered. The characteristic poroelastic relaxation time, *τ_p_*, is on the order of *a^2^*/*D_c_*, where *a* is the radius of the contact area, derived from 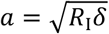 with *δ* the indentation depth, and *D_c_* the cooperative diffusion coefficient(49). *τ_p_* increases with the increase of the tip radius, *R_I_*. For a pyramidal indenter, with a tip radius close to zero, the value of *τ_p_* is small compared to the measurement time, and thus the poroelastic response can be neglected. A spherical indenter is preferable to the pyramidal one for soft samples, because indenting a bigger area prevents penetration of the samples.

We used a spherical indenter with 5 µm tip diameter and the measured Young’s modulus, calculated by using the double-contact model, is comparable with the one of the pyramidal tip, showing that the poroelastic response can be neglected. This calibration procedure was important to show that the obtained results are independent of the indenter geometry and to demonstrate that the double-contact model has to be applied (Figure 2B).

Figure 2C shows that the Young’s modulus increased with increasing monomer concentration, starting from 0.09 ± 0.07 kPa for *C_T_* = 5.9% up to 11 ± 3 kPa for *C_T_* = 11.8%. From within this range, we focused on values between 0.5 – 3.5 kPa which cover most of the values reported of whole cells measured with different techniques(25). The dispersion of elasticity values showed a high variation and the coefficient of variation increased from 20% to 70% with decreasing *C_T_* (inset of Figure 2C). The repeatability of the bead production process and of the elasticity measurements for different batches with three concentrations is shown in Figure S3 (data in Table S2). To understand whether gelation time, combined with production time, affected the final bead elasticity and its variability, droplets were collected at intervals of 5 min in different tubes, and these particles were analyzed by AFM indentation. We observed that bead elasticity varied with collection time as there was a drastic decrease in Young’s modulus after 20 min and no droplet gelation occurred after 25 min (Figure 2D). This behavior may be related to the reactivity of BIS in the pre-gel mixture (cyclization, multiple crosslinking reactions). Also changes in BIS concentration with production time in each droplet(50) could have introduced defects in the polymer network that were responsible for the observed variation in elasticity values. To evaluate the extent of these defects — and to understand the origin of the variability — swelling and elasticity data were used to calculate the structural parameters of the microgels.

### Calculation of structural parameters and poroelastic relaxation time of the beads

Hydrogel structural parameters, such as effective cross-link density, *ν_e_*, strand molecular weight, *M_e_*, and mesh size, *ξ*, are responsible for the microgel response to external mechanical stimuli. Elastic force counteracts the osmotic pressure, and the presence of defects reduces the number of effective elastic chains(51, 52). All these parameters, and the extent of defects as function of *C_T_*, were calculated by combining swelling and elasticity data. The volume fraction of the cross-linked polymer in the equilibrium swollen gel, *v_2_* (inset of Figure 3A)(52, 53) was calculated using the following equation(54):

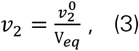

**Figure 3.**
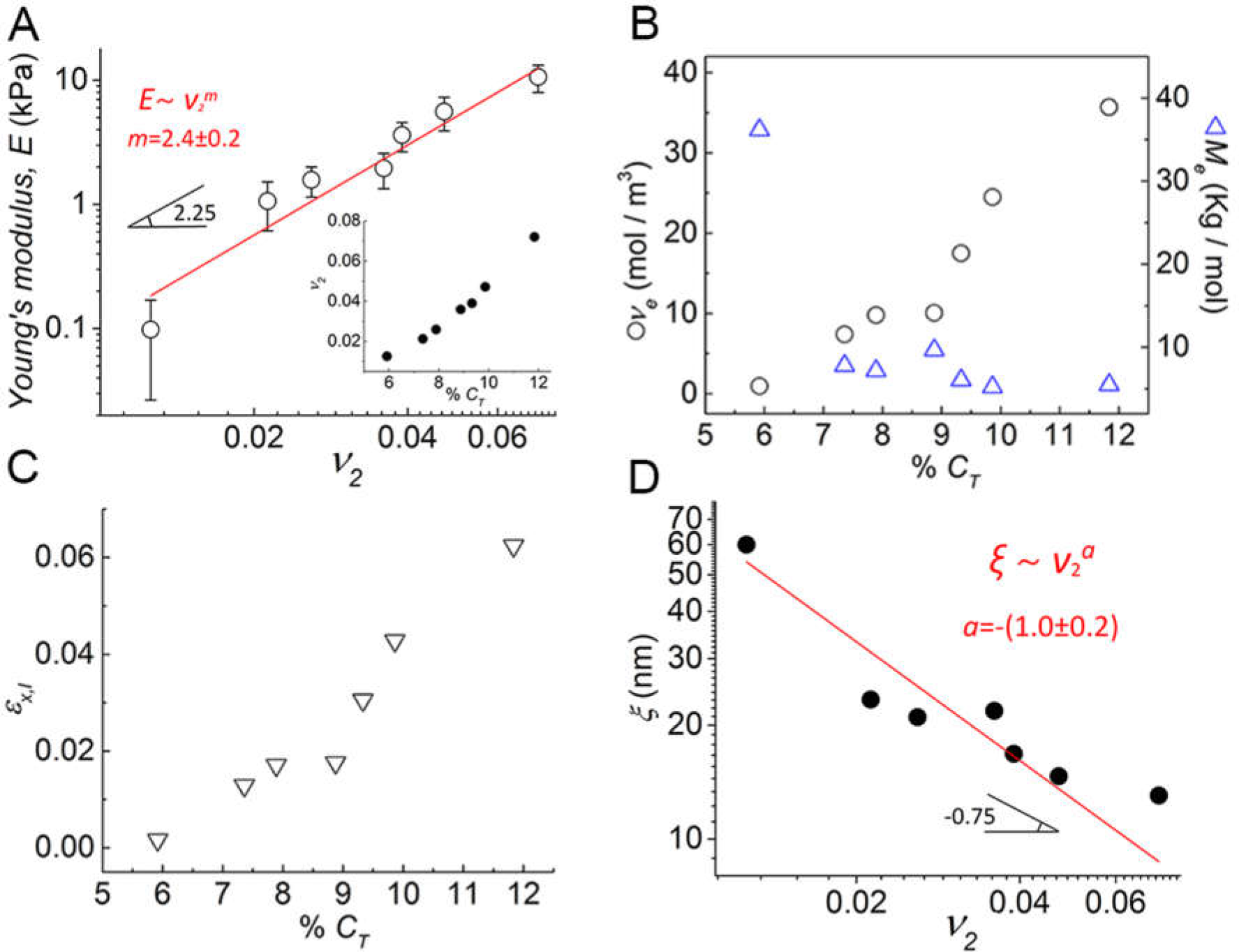
Structural characterization of the microgel beads. (A) Dependence of the bead Youn’s modulus on the volume fraction of the cross-linked polymer in the equilibrium swollen gel (*ν_2_*) in logarithmic scale. The solid red line is the best fit curve by 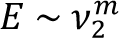 with m = 2.4 ± 0.2. The angle showed the slope of the scaling law for *E* vs. *ν_2_* according to polymer physics (power law 2.25(57)). The inset shows the linear relation between *ν_2_* and *C_T_*. (B) Dependence of the effective cross-linker density, *ν_e_*, and the strand molecular weight, *M_e_*, on *C_T_*. (C) Cross-linking efficiency *εx,l* as a function of *C_T_*. (D) Hydrogel mesh size in dependence on *ν_2_* in logarithmic scale. The solid red line is the best fit curve by 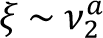 with *a* = – (1.0 ± 0.2). The angle provides a comparison with the slope of the general scaling law for a Gaussian polymer network in a good solvent (power - 0.75)(58).

where 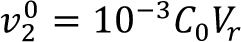 is the volume fraction of cross-linked polymer after gel preparation, *C*_0_ is the initial monomer concentration and *V*_r_ = 52.6 mL/mol is the molar volume of PAAm repeat units(54).

A linear increase in *v_2_* was observed with increasing *CT,* and values varied between 0.013 and 0.070, which are in agreement with previous reports(55). The Young’s modulus *E* was plotted as function of *v_2_* in logarithmic form (Figure 3A) using the relationship between them, based on scaling concepts(56):

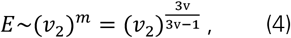

where *v* is the excluded volume exponent that is equal to 3/4 for a good solvent (m = 2.25), 1/2 for a θ solvent (m = 3.0), and 1/3 for poor solvents (m = ∞)(55, 57, 58). Data fitting showed that *v* = 2.4 ± 0.2, indicating a good solvent. Thus, the polymer meshwork was treated as a network of Gaussian chains in a good solvent, where the shear modulus, *G,* of the equilibrated swollen gels is related to the effective cross-linking density *ν*e through the following equation(35, 59, 60):

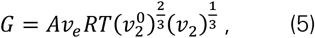

where A = 1–2/f, *R* is the gas constant (8.314 JK^-1^mol^-1^), and *T* the absolute temperature (K). For BIS, the functionality of the cross-linker, *f* = 4. The shear modulus is related to the Young’s modulus by the following equation:

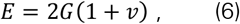

where *ν* is the Poisson’s ratio of PAAm in aqueous medium. Assuming *ν* = 0.5 and combining equations 5 and 6, we get:

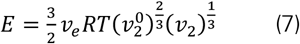

The effective cross-link density, *ν_e_*, was calculated using this equation as all other parameters were known. The strand molecular weight, *M_e_*, (one polymer chain linking two junctions) was calculated as 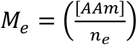, where [AAm] in kg/m³ is the total acrylamide monomer concentration, and 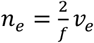 is the number of elastically effective covalent entanglement points(33). The dependence of *ν_e_* and *M_e_* on *C_T_* is depicted in Figure 3B. *ν_e_* increased with increasing *C_T_* from 0.9 to around 36 mol/m³, whereas *M_e_* decreased from 36 to 5 kg/mol. With increasing *CT,* our microgel structure showed a higher number of cross-linking points and shorter connections between them, and this was consistent with the observed density and elasticity increase, as well as swelling decrease with increasing *C_T_*. Cross-linking efficiency was also calculated as *εx,l* = *ν_e_* / *νe,theo*, where 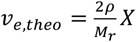 *X* is the chemical cross-link density obtained in *M _r_* an ideal cross-linking process. *M_r_* = 71 g/mol is the molecular weight of repeat units, *X* = (mole BIS / mole AAm) is the cross-linker ratio, and *ρ* = 1.25 g/ml is polymer density(54, 61). The cross-linking efficiency increased with an increase in *C_T_* from 0.002 to 0.062 (Figure 3C), demonstrating that only a certain percentage of the BIS molecules actively participated in the formation of elastic chains.

Microgel porosity, which is important to understand poroelastic effects, was evaluated by calculating the mesh size *ξ* of the polymer network using the following equation(51):

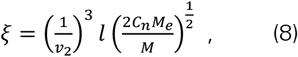

where *M* is acrylamide molecular weight, *C_n_* = 8.5 for PAAm(62) is the rigidity coefficient or Flory characteristic ratio, and *l* = 0.154 nm is the C-C bond length. The variation in polymer mesh size with *ν_2_* could be well described as a power law relation with an exponent *a* = –1.0 ± 0.2 (Figure 3D), which is slightly higher than the value of – 0.75 that is valid for a Gaussian polymer network in a good solvent(58). The correct value of the exponent is still under debate, as another relationship has also been reported in literature, and it is still not clear if a unique law can be used or if various relationships have to be considered depending on the structural properties of the investigated polymer(62). Dolega et al. have reported a pore size smaller than ∼ 10 nm for PAAm microbeads with *E* = 15 kPa(63). This result correlates with our findings, where mesh size decreased with increasing *C_T_* from 60 to 13 nm, with *ξ* ∼ 13 nm for *E* ∼ 11 kPa. According to the calculated pore size, these microgels can be considered as micro-porous, where solute transport is a combination of molecular diffusion and convection in the water-filled pores(64). The cooperative diffusion coefficient(65) *D_c_* was calculated using the following equation, *D_c_* = *kBT*/6*πηξ_h_* (66, 67), where *k_B_* is the Boltzmann constant (1.38×10^−23^ JK^−1^), *T* is the absolute temperature (K), *η* is the solvent viscosity, which is assumed to be equal to that of water (0.91×10^−3^ Pa·s) for PAAm gels(49, 68), and *ξ_h_* is the hydrodynamic screening length, similar to pore size of the gels(67). *D_c_* increased with decreasing *C_T_*, and varied between 0.4×10^−11^ m^2^/s and 1.8×10^−11^ m^2^/s which is in agreement with values of 1 – 5 ×10^−11^ m^2^/s previously obtained for PAAm hydrogels(67). The characteristic poroelastic relaxation time of an indentation experiment is on the order of *a^2^*/*D_c_*, where *a* is the radius of the contact area(67). For a spherical indenter radius of *R_I_* = 2.5 µm and indentation depth *δ =* 1 µm, the radius of the contact area, 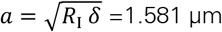(49). According to our calculations, the characteristic poroelastic relaxation time, *τ_p_*, decreased with increasing *CT,* and ranged between 0.14 – 0.63 s. This is comparable to values reported by Kalcioglu et al. in microscale load relaxation experiments(49). The calculated poroelastic relaxation time was also in good agreement with the experimental value obtained from AFM stress relaxation experiments when using a spherical indenter (*τ_p_* = (0.34 ± 0.18) s; Figure 4A). For a pyramidal indenter, no relaxation was observed (Figure 4A). Apparently, the different sample-probe contact areas can affect the variation of the beads’ volume during the indentation process, where a different amount of fluid can leave the network. More specifically, the stress relaxation data provided an evaluation of the beads’ compressibility, through the calculation of their Poisson’s ratio, as 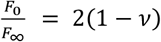, where *F*_0_ is the initial force and *F_∞_* the relaxed force (49). For the pyramidal indenter *F*_0_ = *F_∞_*, *ν* = 0.5 and the beads appear incompressible. For the spherical indenter *F*_0_ = 3.5 nN and *F*_∞_ = 3.0 nN, with *ν* = 0.443 ± 0.007 (Figure 4A), so that for more global stresses the beads are almost incompressible.

**Figure 4.**
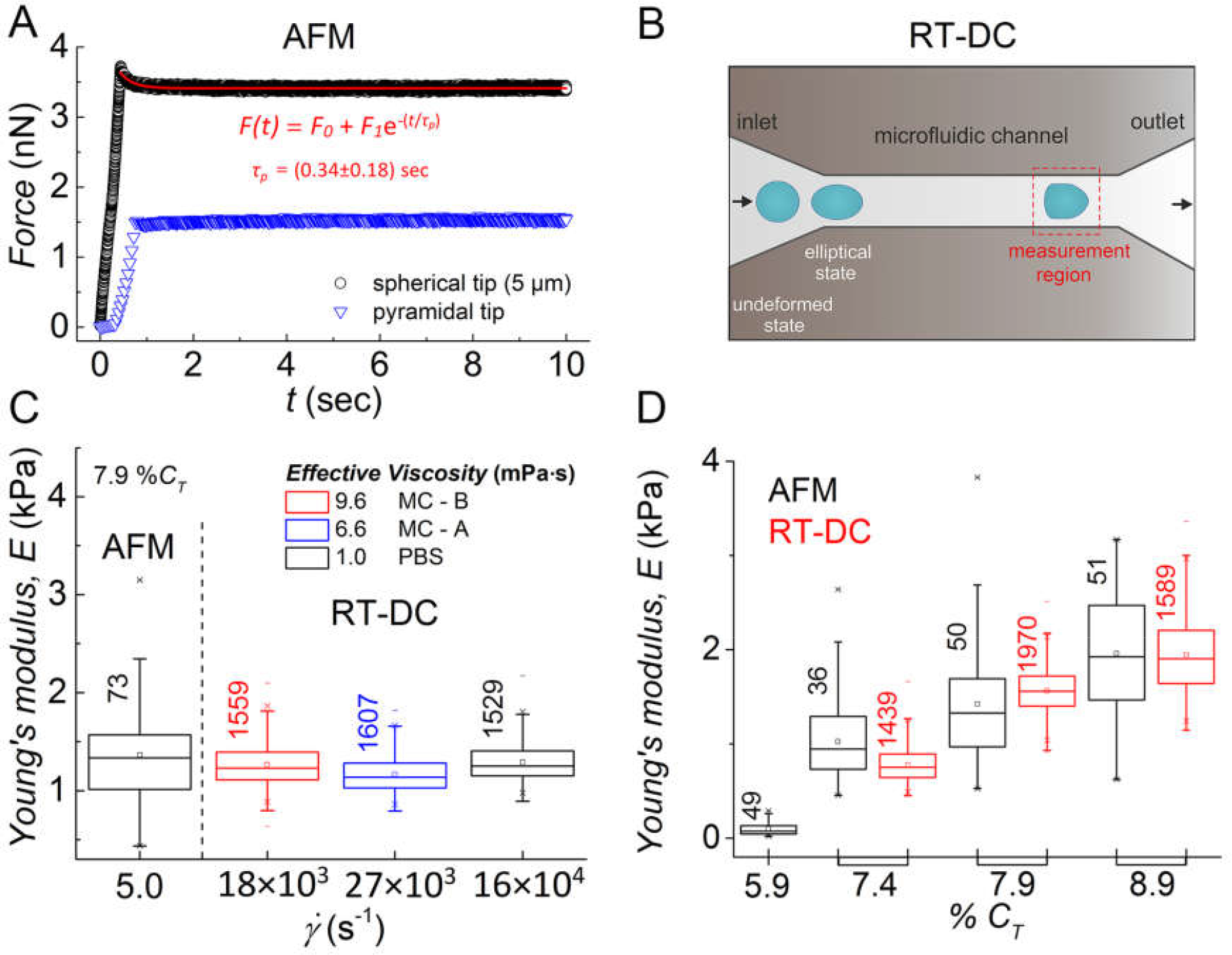
Bead response to deformation applied at different timescales. (A) Representative AFM indentation stress relaxation data on a bead (*C_T_* = 7.9%) using a spherical (black circles) and a pyramidal indenter (blue triangles). Relaxation data with spherical indenter were fitted by using an exponential decay (red solid line), providing the experimental evaluation of the poroelastic characteristic relaxation time *τ_p_ =* (0.34 ± 0.18) s. Indentation data for the pyramidal indenter showed no relaxation. (B) Schematic of the RT-DC setup. The un-deformed beads enter the narrow microfluidic channel assuming an elliptical shape. The beads were measured at the end of the channel (bullet-like shape). (C) Comparison of Youn’s modulus (*C_T_* = 7.9%) determined by AFM and RT-DC at different shear rates, by using different buffers with a viscosity of 9.6 mPa s (red box), 6.6 mPa s (blue box), and 1.0 mPa s (black box). (D) Youn’s modulus obtained with AFM indentation (black boxes) and RT-DC (red boxes) for different gel composition. For *C_T_* = 5.9% only AFM measurement result is shown as beads cannot be detected by RT-DC.

### RT-DC analysis of bead elasticity

In addition to AFM as the standard method in cell mechanical measurements, hydrogel bead elasticity was also investigated using real-time deformability cytometry (RT-DC)(23), to see if at shorter timescales their mechanical response to external stimuli is different from the one observed by AFM. In RT-DC, cells and other spherical objects are deformed by hydrodynamic stresses as they are flowed through a narrow microfluidic channel with a velocity of about 10 cm/s (Figure 4B)(26, 69). Bead deformation *d*, defined as 1 – circularity, was analyzed during the passage through the narrow channel and asymptotically reached an apparently constant value after about 0.5 ms (Figure S4). An exponential fit to the deformation data along the channel provided a characteristic relaxation time, *τ_v_*, of 0.12 ± 0.02 ms (Figure S4). To check if the non-Newtonian behavior and the concentration of the methylcellulose (MC) solution typically used as measurement buffer(69) was affecting our results, multiple measurements were performed using different buffers such as 1x PBS, and MC solutions of different concentration (MC-A, MC-B). The flow rate for each measurement was selected such that most of the beads deformed in the range between *d* = 0.005 – 0.05, to stay within the applicable range of the numerical model used to calculate their Young’s modulus (Figure S5 A,B,C)(26). The effective viscosity and the applied shear rate for the various buffers were calculated using the following equations(70):

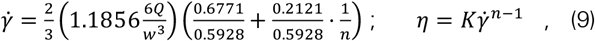

where *Q* is the applied flow rate, *w* is the width of the channel, and the parameters *K* and *n* have been reported previously for MC-A (n = 0.677, K = 0.179) and MC-B (n = 0.634, K = 0.36)(69). The shear rate for AFM indentation was calculated as the ratio between the indentation rate (5 µm/s) and the indentation depth *δ* = 1µm. The data reported in Figure 4C for beads with *C_T_* = 7.9% demonstrated that the measured Young’s modulus was not dependent on MC concentration, liquid effective viscosity, or applied shear rate. Figure 4D shows that the Young’s moduli determined by AFM and RT-DC are comparable. More data about the comparison between AFM and RT-DC analysis for different batches are reported in Figure S6 and Table S3. The consistency of the RT-DC elasticity measurements of six different measurements done on the same batch during two different days is shown in Figure S7 and Table S4. The hydrogel beads made using *C_T_* = 5.9% were only measurable with AFM, as their low refractive index led to insufficient contrast to be detectable in RT-DC. Our findings demonstrate that the cell mechanical-mimics respond as a purely elastic material to applied external stimuli in a wide range of time scales (ms – s). This result ensures that these beads can be used as standard samples for calibration and validation of mechanical measurements at cellular dimensions. Although our analysis of hundreds of thousands of beads demonstrated the consistency of bead production and measurement, high dispersion in elasticity was observed, despite the same nominal composition. To reduce this dispersion, we exploited the correlation between their optical and mechanical properties and used flow cytometry forward scattering (FSC) based sorting to separate the initial bead population into three different sub-populations.

### Flow cytometry forward scattering (FSC) based sorting

Even though the PAAm beads were uniform in size (C.V. < 5.5%), they showed great variation in elasticity that was dependent on their composition (C.V. ∼ 20 – 70%), (inset of Figure 2C). An increase in refractive index was also observed with an increase in elasticity (Figure 5A). The correlation between these two parameters was used to reduce the variability of beads in a given sample by sorting according to their optical properties. In particular, beads with different stiffness showed different FSC signals in flow cytometry. The inset in Figure 5A shows a Gaussian distribution of the FSC signal for beads having the same composition (*C_T_* = 7.9%). The hydrogel particles of a particular batch were sorted into three different populations according to their scattering intensity into “low”, “mid”, and “high” groups. Sorted beads from each population were analyzed by AFM and compared with AFM and RT-DC data of unsorted beads (Figure 5B). The three populations showed different elasticity values each with lower variation than the unsorted population (Table S5). The lowest *C.V.* value (23%) was obtained for the beads sorted in the “mid” group. It became also evident that RT-DC primarily measures stiffer beads from the “mid” and “high” groups, because the more compliant beads with lower contrast might not be detected. With this observation as proof-of-concept, we were able to show that the variation in elasticity of the cell mechanical-mimic beads can be reduced by simply using FSC-based sorting in flow cytometry.

**Figure 5.**
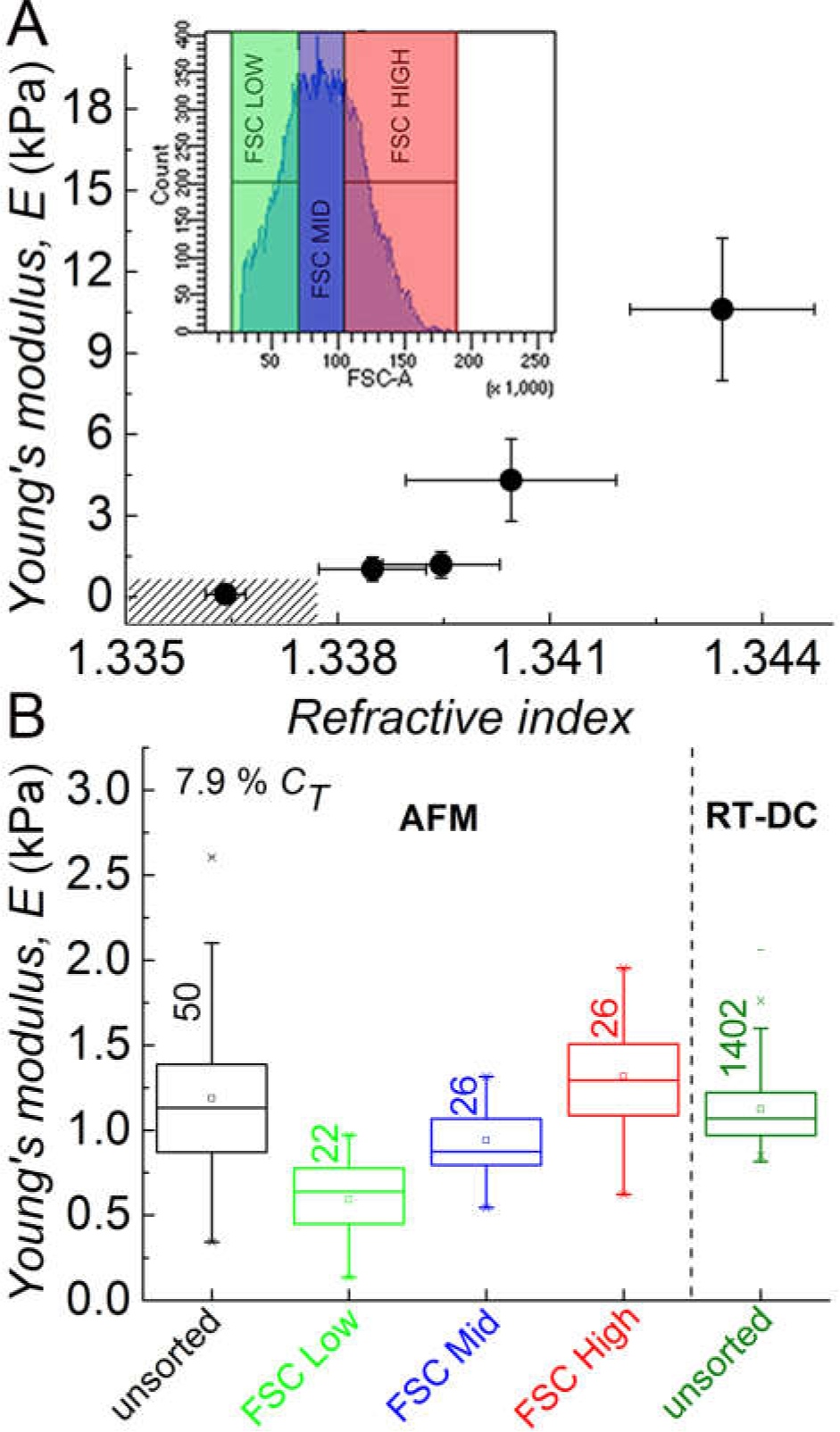
Reduction of bead elasticity polydispersity by FSC sorting. (A) Youn’s modulus as a function of the refractive index of the beads, obtained combining the data reported in Figure 1E and Figure 2C. The dashed area shows the region not detectable by RT-DC. Inset: the graph shows the forward scattering (FSC) signal of the PAAm beads (*C_T_* = 7.9%) by flow cytometry and the FSC sorted regions: low (green), mid (blue), and high FSC intensities. (B) Bead Youn’s modulus, for *C_T_* = 7.9%, after FSC sorting in the 3 subpopulations, measured by AFM indentation, compared with the values obtained on the unsorted beads by RT-DC and AFM indentation.

### Beads functionalized with proteins as passive stress sensors

Our findings demonstrate that PAAm beads can be used as elastic cell mechanical-mimics to calibrate and validate mechanical measurement techniques that are based on the application of controlled deformation or force. Another application is to use these well-characterized microgel particles as stress sensors within cell clusters, either *in vitro* or *in vivo,* to monitor the ability of a cell to generate forces or to measure stress distribution in tissues(71–73). PAAm is chemically inert and the functionalization of the beads with biologically active molecules is crucial in making them cell-interactive(74). PAAm can be functionalized either during or after gelation using sulfo-SANPAH(75) or NHS-ester, respectively(76). We used the NHS-ester, because the production of the NHS-modified hydrogel beads was both faster and easier compared to sulfo-SANPAH. The NHS-ester was added to the oil phase during droplet production and its diffusion in the pre-gel microreactors allowed the inclusion of NHS moieties in the final microgel volume. This method was used to modify the composition of the beads with *C_T_* = 7.9%, but can be applied to the whole range of the total monomer concentration tested. Addition of any amine-containing ligand to the polymerized gel displaces the NHS and results in a covalent bond between the gel and ligand (here, PLL-Cy3)(76). PLL-Cy3 solution was added to the bead suspension and led to a concentration-dependent reaction at the surface, but also inside the bulk of the beads. The fluorescence confocal images acquired in the central plane of the particles showed that functionalization started with small protein clusters on the hydrogel bead surface at low protein concentration (*C_PLL_*) of 0.026 pg/bead. Complete protein incorporation was obtained at *C_PLL_* = 28 pg/bead (Figure 6A). The NHS-modified beads had a Young’s modulus of *E_0_* = 1.6 ± 0.6 kPa. Beads without NHS were produced and used as control, and exhibited a Young’s modulus of 1.4 ± 0.6 kPa. This value is slightly lower than *E_0_*, and it is probably related to intrinsic variations in the microgels (Figure S2-B) rather than an increase in cross-link density due to the addition of NHS. Then the Young’s moduli of the PLL functionalized beads were measured at different concentrations of *C_PLL_*, which led to slightly increased values compared to controls that did not depend strongly on the actual PLL concentration used (Figure 6B). In order to show that the functionalization is effective and that our microgel particles can be used as stress sensors in mechanobiology applications, the beads with *C_PLL_* = 28 pg/bead were embedded in multicellular aggregates of human telomerase reverse transcriptase (hTERT) immortalized human mesenchymal stromal cells (MSC) (Figure 6C). Confocal imaging confirmed stable incorporation of the beads, and cell-induced shape deformations demonstrated the potential of using these PLL functionalized PAAm beads as passive stress sensors.

**Figure 6.**
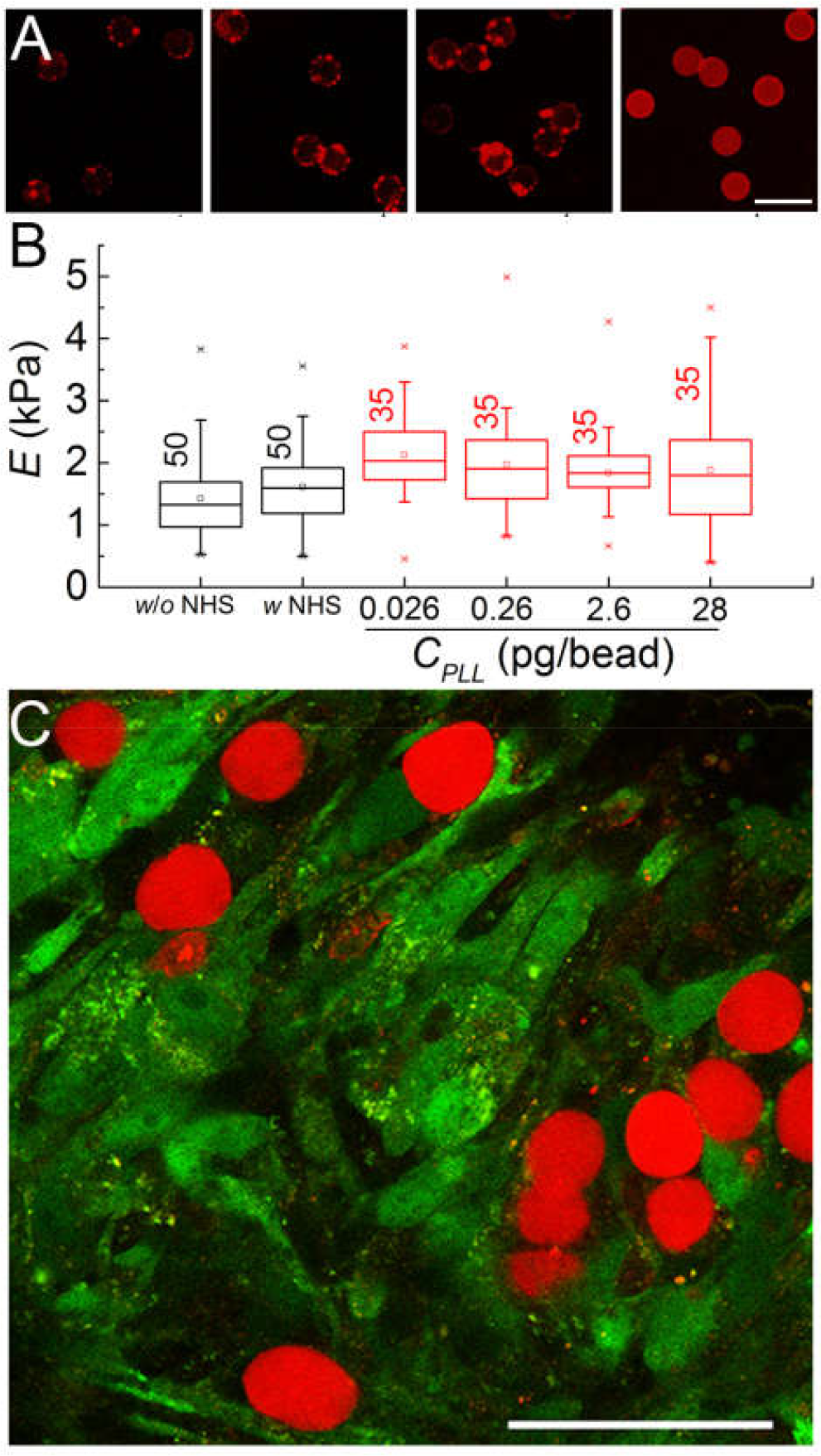
**Beads as cell-scale mechanical probes (functionalization, characterization and application)** (A) Fluorescence images of the beads captured in confocal laser scanning microscopy. PAAm beads were functionalized with Poly-L-Lysine (PLL)-Cy3. Images show PAAm beads with increasing protein concentration (*C_PLL_*) from 0.026 pg/bead (left) to 28 pg/bead (right). Scale bar: 30 µm. (B) Young’s moduli of the beads with the same monomer concentration (*C_T_* = 7.9%) without NHS, with NHS, and after their functionalization with PLL-Cy3 at different PLL concentrations (*C_PLL_*). (C) Fluorescence laser scanning microscopy image of PAAm beads functionalized with Poly-L-Lysine (PLL)-Cy3 in multicellular aggregates of MSCs after a cultivation time of 24 h. Deformations away from spherical shape are clearly detectable. Scale bar: 50 µm.

## DISCUSSION

We successfully demonstrated the high-throughput production of PAAm microgel beads with size and elasticity comparable to that of cells and we verified their homogeneity and purely elastic behavior through an extensive analysis of their optical properties, swelling behavior, and elasticity. The analysis demonstrated that our beads can be used as mechanical probes for validation and calibration of mechanical measurements, and as passive sensors for the analysis of stresses present in biological samples. Previously, hydrogel beads had been produced paying attention only to their size and elasticity. Agarose beads were used as calibration probe, but their analysis by using different techniques (AFM vs. RT-DC) provided different results(24, 25). This discrepancy likely resulted from the viscoelastic mechanical properties of the material(29), which had been interrogated at different timescales with the two techniques. Polyacrylamide is accepted as an elastic material, and microgel beads with specific elasticity have already been produced(63, 77). However, the beads obtained by emulsification showed high size polydispersity. Furthermore, their homogeneity was not demonstrated and the impact of their structural properties on the Young’s modulus and response to different mechanical stimuli was not investigated. The analysis of these aspects is fundamental, especially for polyacrylamide, as different concentrations of monomer and cross-linker lead to the formation of different polymer meshworks, resulting in different physical properties of the hydrogel. It has been demonstrated that hydrogels present topological defects within these meshworks, which affect both their mechanical and structural properties(30, 46, 78). These properties have been investigated for bulk PAAm hydrogel, but not previously for PAAm microgel beads.

### Rational design of PAAm microgel beads

Here, we produced different PAAm microgel beads through the free radical polymerization of a pre-gel solution in a droplet microreactor, keeping the ratio of the cross-linking agent to monomer constant and changing the total monomer concentration. The use of a flow focusing microfluidic device allowed the high-frequency production of monodisperse droplets, eliminating possible variation in the microgel structures due to size variations of the microreactor. To reduce the extent of defects in the final gel, we fixed, based on the phase diagram of polyacrylamide, the ratio of the cross-linking agent to monomer (*C*) at 3.25% and the gelation temperature at 65 °C(79, 80). Studies have already shown that the amount of network defects is also dependent on the total monomer concentration (*C_T_*). Increasing *C_T_* for a fixed *C* decreases the probability of forming network defects(55). Fewer defects provide more elastic chains that counteract the volume increase due to water imbibition. This behavior was also observed for our PAAm beads, where the normalized volume of equilibrium of the swollen gel (*V_eq_*) decreased with the increase of *C_T_* from 5.9 to 11.8% (Figure 1B). The final beads were characterized as a polymer meshwork including topological defects with different amounts of solvent and elastic chains, which in turn affected their optical properties.

### Bead homogeneity

RI analysis was used to evaluate this variation, and at the same time it demonstrated that our beads can be treated as an isotropic material at the microscale (Figure 1E). An increase in RI was observed with increasing *C_T_*. The RI ranged between from 1.3358 to 1.3405, which is close to water (RI of 1.333). For a fixed bead composition, a small variation in the RI was observed, independent of the final bead diameter. The quantitative phase shift analysis used to determine the RI can only provide information about bead uniformity at the microscale, but is not able to resolve density variations at the nanoscale, as already observed by small-angle x-ray scattering measurements(46). However, the constant value of the RI observed along the radial direction (Figure 1G) ensured that TEMED diffusion in the gelling droplets did not vary the cross-link density from the surface to the center of the gel particles. This ruled out a core-shell geometry, which had been discussed before in the literature for bigger gel spheres(40). The homogeneity of the beads was further demonstrated by confocal Brillouin microscopy, where Brillouin shift is not varying within a single bead. The detectable variation of the Brillouin shift is equal to the random error of the measurement system (Figure 1H). This is, to our knowledge, the first direct demonstration of the internal homogeneity of the density and mechanical properties of hydrogel beads. A variation in Brillouin shift between different beads of up to 41 MHz was observed (Figure 1I), which was four times higher than the systematic error of the system of 10 MHz. These variations between different beads, despite the same nominal composition, together with the swelling behavior and the variation of the RI along the gel composition, provided a first evidence of the differences in the mechanical and structural properties of the produced microgels.

### Dispersion of bead elasticity and structural parameters

To obtain more information about the differences in mechanical properties, the bead elasticity was investigated by AFM indentation. The Young’s modulus was shown to increase with increasing *C_T_*, varying between 0.02 and 15 kPa (Figure 2C). Furthermore, although being monodisperse in size (C.V. < 5.5%) and having a low variation in RI (Figure 1E) for a fixed composition, microbeads were dispersed in terms of elasticity as suggested by Brillouin microscopy, with a coefficient of variation that was decreasing with the increase of *C_T_* (*C.V.* = 20 – 70%) (Figure 2C). This suggests that elastic polydispersity is related to the extent of defects, as the probability of defect formation decreases with increasing *C_T_*(55). Furthermore, also the production method can play a role in bead polydispersity for a fixed pre-gel mixture. The droplets were produced starting from 100 µl of AAm, BIS, and APS pre-gel solution, which was consumed in about 30 min. During this period, the co-monomers in presence of the initiator start to interact in the pre-gel volume (pre-network formation)(33, 39). Living radicals are more reactive with BIS than with AAm, leading to the formation of free microgel particles (cross-linked cluster)(46, 81) in the solution during droplet production. As a consequence, the concentration of BIS in the pre-gel mixture and in each droplet was not constant with time, resulting in a variation in composition and structure of each hydrogel bead for a fixed *C_T_*. This behavior was confirmed by AFM indentation analysis of the beads (*C_T_* = 7.9%). For this analysis, the formed droplets were collected each 5 min in different tubes during the entire production process (Figure 2D) and polymerized afterwards separately. A variation of Young’s modulus with production time was observed. Its value drastically decreased after 20 min, and no gelation occurred in the droplets collected during the last 5 minutes. Combining the swelling and the elasticity data by using the phantom network model for a tetrafunctional Gaussian network in a good solvent(34, 59), structural parameters were calculated (Figure 3). The effective cross-link density (*v_e_*) increased with the increase of *C_T_*, followed by a consequent decrease of the strand molecular weight (*M_e_*) as previously observed in bulk gel(33, 57). The values of the cross-linking efficiency (*ε_x,l_*) confirmed that the number of defects increased with the decrease of the total monomer concentration, and showed that only a small percentage (between 0.2 – 7%) of BIS molecules actively participated in the formation of the elastic network(61). On the one hand, the low content in the effective elastic chains made it possible to reach the low values in *E* characteristic for cells. However, the weak hydrogel structure necessitated further characterization to test whether viscoelastic and poroelastic effects influence the bead response to external mechanical stimuli. To validate the use of our beads as mechanical standard it is important to demonstrate that they have an elastic behavior independent of the measurement technique and timescale.

### Beads as cell-like mechanical standard

When a force is applied to induce deformation, the energy can be dissipated via the rearrangements of polymer chains and/or cross-links (viscoelastic response) and via the migration of solvent molecules through the porous structure of the gel (poroelastic response)(49, 82). The different characteristic relaxation timescales of the deformations induced by AFM (Figure 4A) and RT-DC (Figure S2) can provide information about the relative importance of poroelastic or viscoelastic effects. For AFM indentation of hydrogels, poroelastic relaxation is dominant over the viscoelastic contribution(49). Viscoelastic properties can be investigated by using AFM high-frequency indentation (up to 100 kHz) leading to correspondingly shorter measurement timescales(83). We calculated a theoretical poroelastic characteristic relaxation time of 0.14 – 0.63 s (depending on composition), which is in the range of the applied indentation time (0.2 s) and comparable with that found experimentally (*τ_p_* = 0.34 s) by AFM stress relaxation (Figure 4A). Furthermore, AFM analysis with pyramidal indenter showed an incompressible behavior of the beads (Figure 4A), Poisson’s ratio of 0.5, providing a Young’s modulus comparable to the one obtained by using a spherical indenter (Figure 2B). To demonstrate that poroelastic effects can be neglected at shorter timescales, the AFM results were compared to those obtained by RT-DC.

An analysis of the bead deformation dynamics in RT-DC showed that the deformation reached a (quasi-) steady-state with a characteristic (viscoelastic) relaxation time of about 0.12 ms (Figure S5). The deformation did not vary further within the 3 ms after they entered the narrow channel, at which time they are analyzed in the standard RT-DC measurement. In order to also detect the poroelastic dissipation, RT-DC measurements would have to be done after 0.14 – 0.63 s, which is practically not possible. Furthermore, also the beads’ compressibility can affect the measurements. Several techniques have been used to measure the Poisson’s ratio of PAAm bulk gels, reporting a quite large range of values, *ν* = 0.32 – 0.49(49, 84–88). Apparently, at small scale, beads can behave as compressible or incompressible material depending on the method, deformation time, and probe-size used for their analysis. During RT-DC there is no time for the fluid to leave the network, and beads can be considered incompressible. For AFM indentation we showed that their compressibility is dependent on the geometry of the indenter, with *ν* = 0.5 (pyramidal indenter) and *ν* = 0.44 (spherical indenter). The quasi-incompressible behavior (*ν* = 0.44) observed for spherical indenter is in good agreement with previous results(86–88), and induces only a slightly variation in bead elasticity. Specifically, assuming the beads as incompressible (*ν* = 0.5) we calculated a Young’s modulus value which is about 6% lower compared to the value obtained with *ν* = 0.44.

RT-DC analysis was performed on compliant beads that mimic cell elasticity, for which discrepancy in cell measurements with AFM and RT-DC was observed(25). The good agreement between AFM indentation data and RT-DC data for beads with *C_T_* varying between 7.4 and 8.9% showed that our beads have a mechanical response independent of the measurement technique, with their vastly different measurement timescales, in a range of elasticity from 0.5 to 3.5 kPa (Figure 4D). Our finding was corroborated by further RT-DC results. The Young’s modulus extracted did not vary when changing the measurement buffer from 1x PBS to solutions with different MC concentrations (Figure 4C). This demonstrated that adding MC to the buffer (not present in AFM experiments), as well as its non– Newtonian behavior(69), and performing experiments at different shear rates 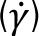 did not affect the bead response to the stresses applied. The remaining difference in bead elasticity dispersion between RT-DC and AFM could be related to the difference in focus and contrast between measurements, which affect their contour detection in the microfluidic channel during RT-DC measurement and adds a random error. Other random errors in the AFM measurements, potentially related to the use of different cantilevers, variations in the colloidal probe, or the degree of bead adhesion to the substrate, might also contribute. There is an additional systematic bias in the RT-DC measurements that becomes very obvious for low concentration beads. It was not possible to analyze beads with *C_T_* = 5.9% by RT-DC. Combining this finding with the relation between Young’s modulus and RI (Figure 5A), we concluded that beads with RI < 1.3376 (corresponding to Young’s modulus, *E* < 0.7 kPa) cannot be detected by RT-DC.

### Reducing bead polydispersity in elasticity through the relation between their mechanical and optical properties

Our results showed that bead swelling and elasticity respectively decreases and increases with the increase of *C_T_* (Figure 1B, 2C), and that the probability of defects decreases with increasing *C_T_* (Figure 3C). This generated a variation in bead density correlated to their variation in elasticity, where beads with a fixed value of *C_T_* provided different FSC values during flow cytometry measurement (Figure 5A inset). This behavior was used to reduce bead polydispersity in stiffness by sorting them with flow cytometry and subsequently measuring their stiffness with AFM. The sorting demonstrated that the overall bead population detected by RT-DC can be reconstructed considering the beads collected in the “high” and “mid” region of the FSC curve (Figure 5B). This confirmed that RT-DC mainly detected stiff beads and missed the compliant ones, because of their low contrast. More importantly, sorting of PAAm beads based on their FSC signal in a conventional flow cytometer can be used as a practical solution to reduce the variability in elasticity in the sorted populations. This has important implications for the production of beads with different methods, which might otherwise have unacceptable heterogeneity in elasticity.

### Beads as passive cell-scale stress sensors

Further functionality was added to the beads to enlarge their applicability to the *in vivo* measurements of stresses present in cell clusters or tissue. We demonstrated a simple and fast method to introduce functional groups into the gel during droplet production. The Young’s modulus of the NHS-modified microgel beads produced using *C_T_* = 7.9% had a value of 1.6 ± 0.4 kPa (Figure 6B). The addition of PLL to the polymerized microgel introduced charges that could, in turn, affect bead elasticity. We measured higher elasticity for the beads functionalized with PLL, probably due to the formation of multiplets in the gel acting as additional cross-link points(53). The potential of the uniformly coated beads (with *C_PLL_* = 28 pg/bead) as cell-scale sensors was tested by their incorporation in multicellular aggregates of human hTERT-immortalized human MSCs (Figure 6C). The introduction of fluorescent PLL into the bulk of the beads promoted cellular adhesion and facilitated fluorescent confocal imaging. This offers the opportunity to use such functionalized beads as probes to quantify mechanical stress acting on them at the cellular scale. While molecular scale stress sensors are now frequently used, most commonly in the form of FRET dyes attached to calibrated molecules that unfold when being part of a molecular force chain, cell-scale stress sensors are still not used widely(89). Sensors based on oil droplets have been used to measure stresses in tissue, but with such incompressible liquids isotropic and shear stresses cannot be quantified(71, 72). Recently Dolega et al. showed that isotropic stress can be measured by using PAAm beads(63), but in this case external compressive stress was applied to multicellular spheroids by using osmotic pressure. The ability to characterize the material structure and mechanical properties of our beads in detail and to adjust their properties to mimic cells specifically, represents a unique way to measure both isotropic and anisotropic, as well as shear stresses passively when the beads interact with individual cells, such as during migration or phagocytosis, or inside cell clusters, organoids, or tissues.

## CONCLUSION

In this work, we demonstrated the reliable high-throughput production of elastic cell mechanical-mimics, and their suitability to act as calibration standard for different mechanical techniques with vastly different measurement timescales. Refractive index tomography and Brillouin microscopy confirmed for the first time explicitly the internal homogeneity of the beads produced. The remaining variability in elasticity can be ascribed to unavoidable defects in the polymer network and different relative concentrations of cross-linker. This variability can be reduced by sorting of populations defined by their forward scattering behavior using conventional flow cytometry. Overall, the results demonstrate that these microgel beads are a powerful tool for the cross-comparison of results obtained from different cell mechanical characterization techniques, and for the validation of upcoming novel techniques. Finally, we illustrated a simple and flexible way to functionalize the elastic cell mechanical-mimics with proteins for further applications, such as measurement of stresses present when interacting with individual cells, in organoids, or in living tissue during development. This opens new perspectives for the analysis of stresses exerted not only at the cellular scale, but also inside cells, for example in synthetic biology, where beads functionalized with DNA can be used as artificial nuclei to understand stresses acting on nuclei during cell division. Many other areas of application in biophysics, biology, and biomedicine are evident.

## ACKNOWLEDGMENTS

We thank the Microstructure Facility and the Flow Cytometry Facility of the CMCB (both in part funded by the State of Saxony and the European Fund for Regional Development — EFRE), respectively, for the production of the microfluidic devices and microgel beads, and for the bead sorting. We thank Yara Alsaadawi for help in bead production and Florian Oltsch for writing the FIJI macro used for the size distribution analysis. We thank Dr. Mikhail Malanin for IR analysis of the surfactant, and Kathrin Pöschel for CMC measurements of the surfactant. We also thank Matthias Schieker for providing the SCP-1 cell line. We thank Jonas Tegenfeldt, Jason Beech, Kushagr Punyani, and colleagues from Lund University and other members of the LAPASO ITN for fruitful discussions. This project has received funding from the European Research Council Starting Grant “LightTouch” (grant agreement number 282060 to J.G.) and from the Alexander-von-Humboldt Stiftung (Humboldt-Professorship to J.G.). J.T. received funding from the Federal Ministry of Education and Research (BMBF, “Biotechnologie2020+Strukturvorhaben: Leibniz Research Cluster, 031A360C), and the German Research Foundation (DFG, TH 2037/1-1), which is gratefully acknowledged.

## MATERIALS AND METHODS

### Materials

Acrylamide suitable for electrophoresis ≥ 99%, N,N′-Methylenebis acrylamide (Bis-acrylamide) 99%, N, N, N′, N′-Tetramethylethylenediamine (TEMED) ∼ 99%, acrylic acid N-hydroxysuccinimide ester ≥ 90%, 1H,1H,2H,2H-Perfluoro-1-octanol (PFO) 97% and Span® 80 were purchased from Sigma-Aldrich Chemie GmbH, Schnelldorf, Germany. Ammonium Persulphate was obtained from GE Healthcare, Germany. HFE 7500 was purchased from Ionic Liquids Technology, Germany. AZ 15n XT photoresist was purchased from MicroChemicals GmbH, Ulm, Germany. Polydimethylsiloxane (Dow Corning Sylgard® 184) was purchased from Ellsworth Adhesives, Glasow, UK. Aquapel® was obtained from Pittsburgh Glass Works LLC, Pittsburgh, USA. Poly-L-lysine, Cy3 labeled (∼25kDa, 2 mg in 0.5 ml H2O) was ordered from Nanocs Inc. (www.nanosc.net). Hexane was obtained from Carl Roth, Germany. Tris-Buffer (pH 7.48) was prepared with 10 mM Trizma® base (Sigma-Aldrich Chemie GmbH, Germany), 1mM EDTA (Sigma-Aldrich Chemie GmbH, Germany) and 15 mM NaCl (Merck Chemicals GmbH, Germany) in ultrapure water. 50 mM HEPES solution (pH 8.22) was prepared by dissolving HEPES (PUFFERAN®) ≥ 99.5 % (Carl Roth, Germany) in ultrapure water. Acetonitrile was purchased from Merck Millipore, Methanol from Acros, Ammonium hydroxide solution (28-30 %, based on NH3 content) from Sigma-Aldrich, Krytox® 157 FSH from DuPont, and HFE 7100 from Iolitec. 1x PBS containing no Ca and Mg was used and methylcellulose (MC) solutions, (MC-A = cell buffer; MC-B = cell buffer B), were kindly provided by Zellmechanik Dresden.

## Methods

### Fabrication of microfluidic devices

The master template, with relief structures (height 20 µm) reproducing the device geometry, was realized by a photolithography process (mask aligner EVG620), by using AZ15nXT photoresist, deposited by spin coating (Laurell WS-650Mz-23NPP spin processor) on a 4’’ silicon wafer. The master features were replicated on a polydimethylsiloxane (PDMS) element by replica molding process. Base (Sylgard® 184) and curing agent were mixed in ratio 10:1 (w/w), degassed, poured on the master surface and polymerized in an oven at 75 °C for 1.5 hours. Afterwards, the PDMS replica was peeled off from the master and punched in correspondence of the inlet and outlet chambers by using a biopsy puncher (I.D. 1.5 mm). Finally, the PDMS element was bound on a glass coverslip (40×24 mm^2^), (thickness 2, Hecht, Germany) by the activation of their surfaces with an air plasma treatment (50 W, 30 s, Gambetti, Tucano plasma system). The droplets generator device(38) included two inlet and one outlet chambers and its geometry at the cross flow region was characterized by two perpendicular channels with a width of 15 µm and a height of 20 µm. The RT-DC microfluidic device(23) included two inlets and one outlet chambers and is characterized by a squared narrow constriction, 20 µm wide and 300 µm long.

### Surfactant synthesis

The ammonium carboxylate salt of Krytox® was synthesized according to the literature with slight modifications(90). Briefly, 10 g Krytox® 157 FSH was dissolved in a mixture of 60 mL methanol and 30 mL HFE 7100. After stirring until full dissolution, 25 mL 0.1 N ammonium hydroxide was added dropwise. The mixture was allowed to react overnight under vigorous stirring. Removal of the solvent under reduced pressure, followed by drying under high vacuum gave a pale, viscous oil(91).

### Preparation of polyacrylamide hydrogel beads

The microfluidic device inner walls were functionalized by flushing Aquapel® inside the microchannels. The solution was removed by blowing the device with an air gun and leaving it in the oven at 65 °C for 10 mins. The oil solutions were prepared, adding to the HFE-7500, ammonium Krytox® surfactant (1.5% w/w) and TEMED (0.4% v/v) for the standard PAAm beads and with the further addition of N-hydroxysuccinimide ester (0.1% w/v) for the NHS-modified beads. Before using, they were filtered through 33 mm syringe filter (hydrophilic PVDF 0.22 µm membrane, Merck Chemicals GmbH, Germany). The pre-gel mixtures were obtained by mixing different amount of acrylamide (AAm), bis-acrylamide (BIS) and ammonium persulphate (APS) stock solutions prepared by dissolving them in 10 mM Tris-buffer with a concentration of 40% w/w, 2% w/w and 0.05% w/v, respectively. The final volume of each of the polyacrylamide solutions was adjusted to 545 µl by adding the Tris-buffer (pH 7.48) to the AAm, BIS, APS mixture. In all the solutions the molar ratio of bis-acrylamide to acrylamide was constant as 1:61.5. Two vials containing respectively 1 ml of oil and 100 µl of polyacrylamide solutions were connected to the inlet chambers of the device via FEP tubing (I.D. 250 µm, O.D. 1.5 mm, Postnova Analytics GmbH, Germany). The flow, through the tubing towards the device, was activated and controlled by pressurizing the liquids inside the vials by using the Fluigent MFCS™-EX microfluidic controller. The controller was equipped with two channels able to provide a maximum pressure of 1000 mbar. The resulting gel mixture in oil emulsion was collected in a 1.5 ml Eppendorf® tube through a FEP tubing (I.D. 250 µm, O.D. 1.5 mm) connected to the outlet chamber of the device. The time, *t_C_*, needed to consume 100 µl of polyacrylamide mix (*V_sol_*) was recorded in order to calculate the droplet frequency. The mean droplet volume, 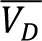, was calculated using the mean value of the droplet diameter in oil 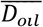, obtained by the analysis of their size distribution. The droplet frequency was calculated by using the following equation: 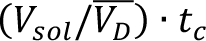. The produced droplets were then incubated overnight at 65 °C to bring them to a complete polymerization. The final hydrogel beads in oil were washed three times via centrifugation (5000 rcf, 1 min) with each of 20% v/v PFO in HFE-7500, 1% v/v Span® 80 in Hexane and 1x PBS (without Mg and Ca) solutions. The final beads suspension in 1x PBS was stored at 4 °C. Phase contrast images of the beads in oil and PBS were acquired under an inverted microscope (Zeiss, Axio Observer.A1) by using a Plan-Apochromat 20x/0.8 Ph2 and analyzed by using a macro implemented on freeware Fiji in order to measure their size distribution. NHS-modified beads were centrifuged (5000 rcf, 1 min) and re-suspended in HEPES solution (pH 8.22) for their functionalization with Poly-L-Lysine (PLL) conjugated with Cy3 fluorophores. 40 µg, 4 µg, 0.4 µg, 0.04 µg of PLL were added to different vials, each containing about 1.5×10^6^ beads. After overnight incubation, the beads were washed three times via centrifugation (5000 rcf, 1 min) with 1x PBS and stored at 4 °C. The functionalized beads were viewed by confocal laser scanning microscopy with a Zeiss LSM 700, objective C-Apochromat 40x 1.2W Korr and 555 nm laser (10 mW).

### Refractive index analysis

A low concentration bead solution was placed in between two coverslips. To ensure the light passes straight through the sample, the two coverslips were arranged parallel to each other by placing double-side tape in between. To determine the RI of the PAA beads, quantitative phase microscopy was performed to measure the phase shift introduced by the beads(92). The quantitative phase images were recorded with the quantitative phase-imaging camera (SID4Bio, Phasics S.A.) with the same setup as described in Schürmann *et al.*(92). In brief, the acquired phase comprises the product of height and average RI at each x-y-position in the image. Height can be estimated assuming a spherical shape of the beads. The homogeneity of the beads was checked by determining the radial refractive index. The radial refractive index profiles were computed from representative phase images by inverting the imaging process. This inversion assumes that the hydrogel beads are rotationally symmetric, i.e. that the phase images look the same for all possible observation angles. The inversion was performed with optical diffraction tomography in the Rytov approximation(42, 43) using the backpropagation algorithm as implemented in the ODTbrain library version 0.1.6(43).

### Confocal Brillouin Microscopy

The Brillouin shift was measured by confocal Brillouin microscopy, using the set-up described in a previous publication(45), which has an optical resolution of below 1 µm in the lateral plane and approximately 3 µm in the axial direction. For the measurement, the beads were placed in a glass bottom petri dish filled with PBS. The Brillouin shift was then measured at a height of eight µm above the coverslip in the center of the beads. For the measurement of the Brillouin shift on a single bead, reported in Fig. 1H, we measured 41 times 41 points with a spatial resolution of 0.75 µm. For the same analysis on 15 different beads (Figure 1I), we measured 100 points within a square area with 7 µm edge length.

### Atomic Force Microscopy (AFM)

All AFM indentation measurements were performed using a Nanowizard I AFM (JPK Instruments) mounted on an inverted optical microscope (Axiovert 200, Zeiss). For presented elasticity measurements standard pyramidal tipped cantilevers (MLCT, nominal spring constant k = 0.03 N/m, Bruker) and cantilevers modified with a spherical tip (Arrow-TL1x20-50, nominal spring constant k = 0.035 – 0.045 N/m, NanoAndMore GmbH) were used. The cantilevers were calibrated by thermal noise method before each experiment(93). Spherical tipped cantilevers were prepared by gluing polystyrene microspheres (diameter: 5 µm, Microparticles GmbH) to the end of the tip-less cantilever using a two-component epoxy glue (Araldite). During the experiments, cantilever tips were aligned over the center of the beads and individual force-distance curves were acquired with 5 µm/s approach and retract velocity and with contact forces ranging from 1 – 8 nN. Forces were chosen to keep the indentation depth constant at 1 µm and Poisson’s ratio was set to 0.5. The Youn’s modulus was extracted from approach force-distance curves using JPK data processing software (JPK Instruments). To prevent motion during indentation measurements the PAAm beads were immobilized on the bottom of a plastic petri dish using a cell adhesive protein solution (CellTak, Cell and Tissue Adhesive, Corning). All measurements were carried out in PBS and at room temperature.

Stress relaxation experiments were performed using a Nanowizard 4 AFM (JPK Instruments) mounted on an inverted optical microscope (Observer D1, Zeiss). Cantilever and sample preparation were the same as for indentation measurements. Individual force-distance curves were acquired with 15 µm/s approach and retract velocity, with a holding time of 10 s, and a contact force of 2.8 nN (4 – 5.4 nN) and indentation depth of 1.5 µm (0.9 – 1.9 µm) for the pyramidal indenter (spherical indenter).

### Real time deformability cytometry (RT-DC)

The RT-DC measurements were carried out by using the same setup and device described in a previous publication(23). Briefly, PAAm beads were suspended at a concentration of 3×10^6^ beads/ml in different buffers, such as 1x PBS, and methylcellulose (MC) solutions of different concentration (MC-A, MC-B). The MC concentration for MC-A and MC-B was respectively 0.5% (w/v) or 0.6% (w/v) methylcellulose in 1x PBS, providing respectively a viscosity of 15 mPa·s and 25 mPa·s. The deformation *d* is defined using circularity 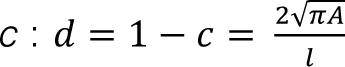 where *A* is the projected area and *l* the perimeter of the object. The lower limit of the measurable deformation is given by the resolution of the camera and it is 0.005. The upper limit is equal to 0.05 to meet the validity range for calculation of the Young’s modulus(26). By adjusting the flow-rate for each sample, we made sure to exert enough force to reach comparable deformations in the range of 0.005 – 0.05. A method that uses numerical simulations, to describe the forces and the shape evolution of elastic objects in microfluidic channels allows calculating the elastic modulus *E* of individual beads(26). Corrections were applied considering the non-Newtonian behavior of the MC solutions, and viscosity variation due to temperature. To disregard all objects with concave parts, we computed the area ratio as shown in a previous publication(94). Shortly, the area of the convex hull of the contour and the area of the original contour are divided by each other. All beads with an area ratio above 1.05 were filtered out and were not considered in further analyses. Additionally, all events below a deformation of 0.005 and above 0.05 were filtered out. All analyses were done using the open source software ShapeOut(95).

### Flow Cytometry

PAAm beads in 1x PBS were analyzed and sorted by flow cytometry based on the forward-scattered light signal. Doublet discrimination was carried out. Sorting boundaries were chosen with regard to the plateau of the signal distribution. All measurements were performed on a BD FACS Aria II SORP (BD Bioscience) according to manufacturer’s instructions. Data analysis was done using BD FACSDiva 8 software.

### Virus particle production and Lentiviral plasmid transduction

The lentiviral transfer vectors were generated as previously described(96). The vector pRRL.PPT.SF.eGFP-T2A-luc2.pre* without any target was kindly provided by Axel Schambach (Hannover, Germany). Lentivirus particles were produced by co-transfection of 293T cells with a VSV-G envelope plasmid, a Gag/Pol plasmid and the lentiviral vector carrying eGFP and firefly luciferace (pRRL.PPT.SF.eGFP-T2A-luc2.pre*). 293T cells were seeded at a density of 65,000 cells/cm^2^ and grown in complete DMEM medium (Gibco, Invitrogen, Carlsbad, CA) with 10% fetal bovine serum (FBS, Gibco, Invitrogen) up to 70% confluency. Cells were then infected with the lentivirus vector mix prepared in DMEM without serum, containing 10 µg/mL Polyethylenimine (Sigma-Aldrich). Medium was replaced after 8 h with complete DMDM and again after another 12 h with complete DMEM. Viral supernatant was harvested 24, 48, and 72 h after transfection, centrifuged at 300 g for 5 min, filtered using 0.45 µm pore size filter (Corning Inc, Corning, NY), and frozen in aliquots at -80 °C. Lentiviral titer was determined using a QuickTiterTM Lentivirus Quantitation Kit (Cell Biolabs, San Diego; CA).

### Transduction of SCP-1

The human telomerase reverse transcriptase (hTERT) immortalized human mesenchymal stromal cell line SCP-1(97) was expanded in complete DMEM medium (Thermo Fisher Scientific, Waltham, MA) with 10% FBS (Gibco, Invitrogen). SCP-1 cells were transduced at 70% confluence. Transductions were performed over night at 37°C with 1:2 diluted virus vector supernatant (7.5 x 10^9^ virus particles/mL) supplemented with 1 mg/mL protamine. Media was replaced after 24 h with complete DMEM. After recovering 24 h in complete DMEM medium, a second transduction was done for 24 h. Transduction efficiency of GFP was confirmed by FACS analysis 72 h after the second transduction.

### Multicellular aggregate preparation

MSCs were maintained in a humidified atmosphere of 5% carbon dioxide and cultured in low-glucose Dulbecco’s modified Eagle medium (DMEM, Gibco) supplemented with 10% FBS (FBS, Gibco). Multicellular aggregate formation was induced according to hanging drop method. Cell suspension was mixed with PAAm beads functionalized with PLL and seeded in form of 70 µl drops of an inverted lid of a 35 mm petri dish for a culture period of 24 h.

### Statistical Analysis

In the box plots the mean is shown as straight line and the boxes are determined by the 25^th^ and 75^th^ percentiles. The crosses are determined by the 1^st^ and 99^th^ percentiles and the whiskers are the minimum and maximum value. Number of measurements (*n*) is given in the respective box in the diagrams, except Figure 2B where always 31 beads were measured. Coefficient of variation (*C.V.*) is defined as ratio of standard deviation to mean value. Data depicted in Figure 1A show mean ± standard deviation (S.D.) of three different batches produced on different days. Data points with error bars in Figure 1B represent mean ± standard deviation (S.D.) (number of measurements (*n*) is given at Table S1). The data and the propagating error reported in the inset in Figure 1B were derived from data reported in Table S1. All data points reported in Figure 3 were derived using data reported in Figure 1 and Figure 2. The value of the poroelastic characteristic relaxation time, *τ_p_,* reported in Figure 4B is given as mean± standard deviation (S.D.), calculated using the data of independent stress relaxation measurements done on 27 beads with *C_T_* = 7.9%.

## SUPPLEMENTARY MATERIAL

**Figure S1.**
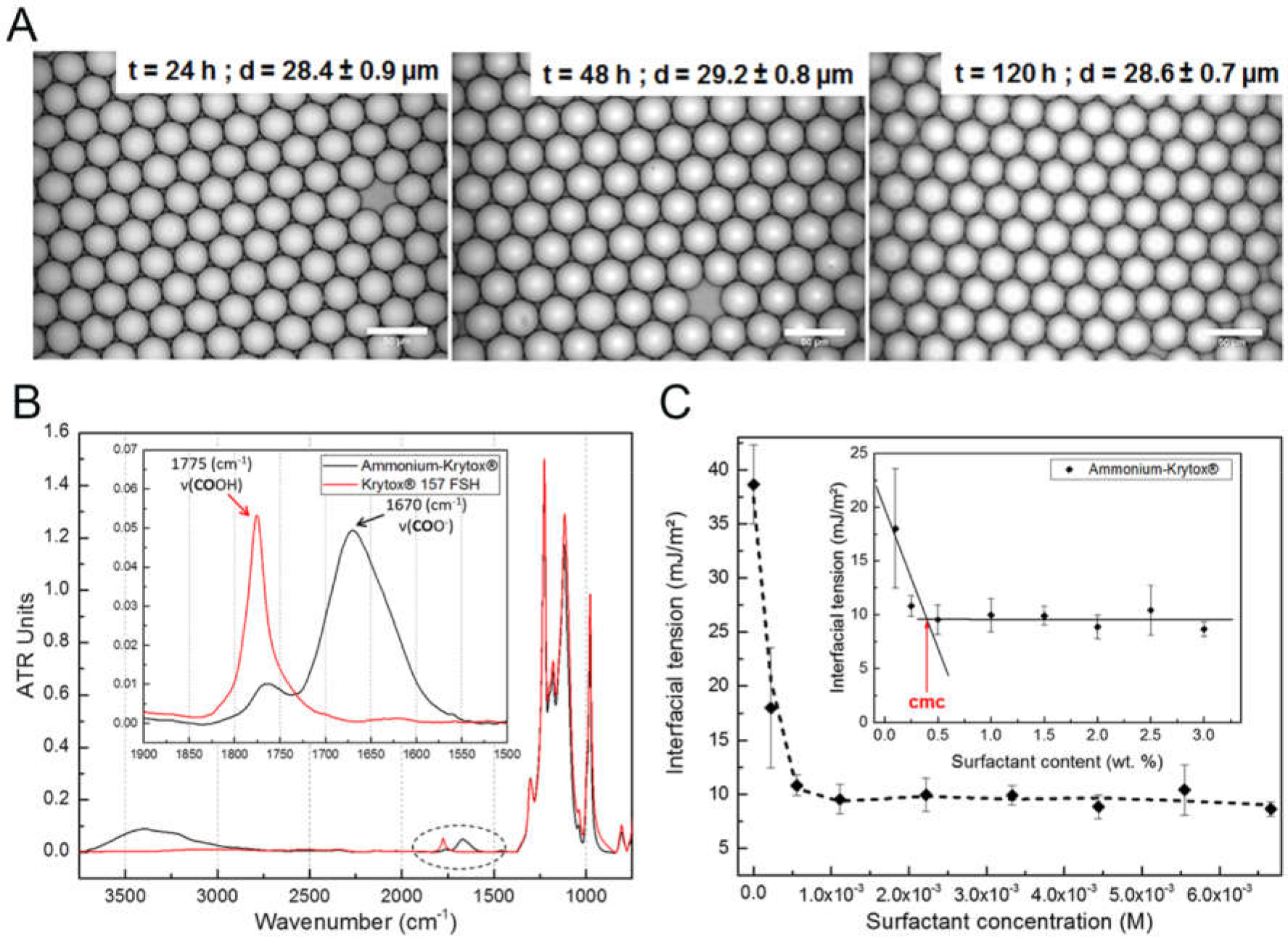
Ammonium Krytox^®^ surfactant, emulsion stability and surfactant analysis. (A) Long-term emulsion stability test. A water-in-fluorinated oil emulsion – stabilized by the ammonium Krytox^®^ surfactant in HFE 7500 – is stored for several days, and its stability confirmed by droplet size distribution analysis (scale bar, 50 µm). (B) FTIR spectrum of the ammonium-Krytox^®^ surfactant and Krytox^®^ 157 FSH as key starting material showing the conversion of Krytox^®^ 157 FSH-carboxylic acid groups (1175 cm^-1^) to ammonium-Krytox^®^ carboxylate (1670 cm^-1^).(C) Determination of the critical micellar concentration for ammonium-Krytox^®^ by pendant droplets analysis. HFE 7500/surfactant solution is extruded from a syringe into a cuvette of double-distilled water.

**Figure S2.**
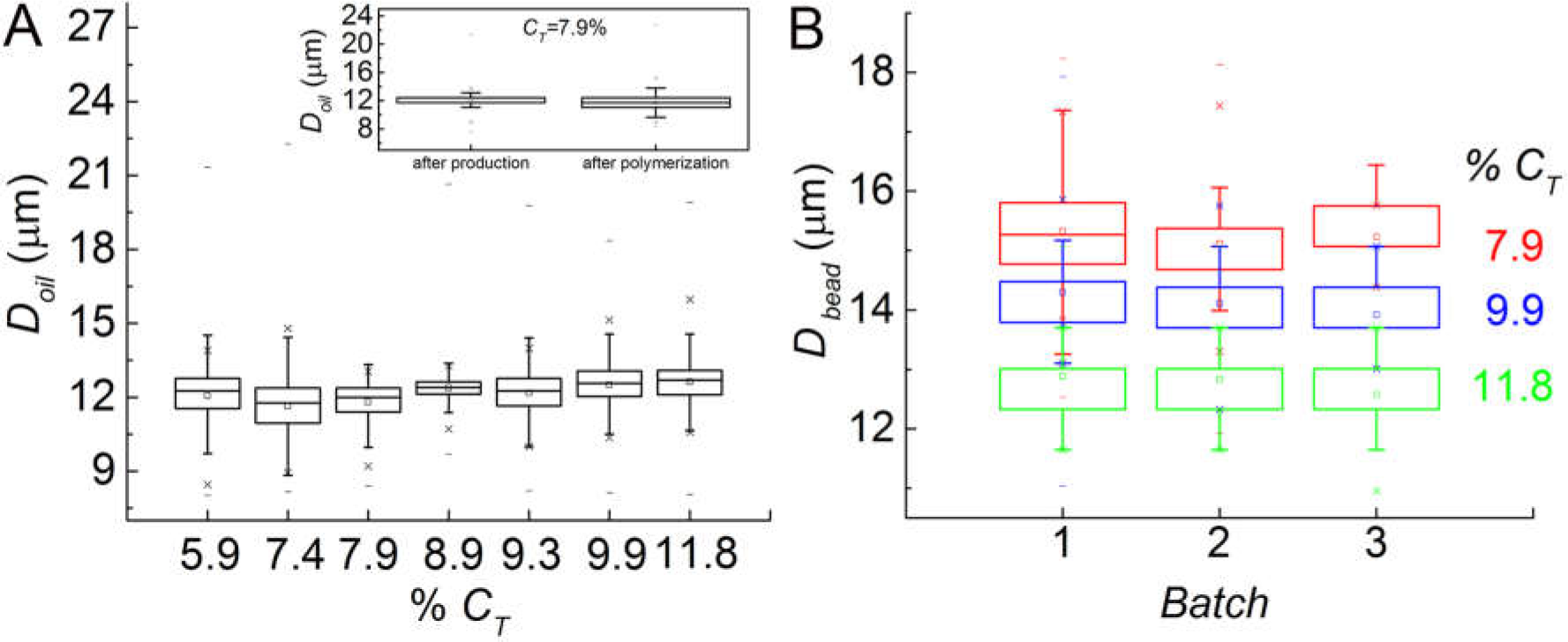
Repeatability of bead swelling behaviour. (A) Distribution of droplet diameter for different monomer concentrations (*C_T_*) measured with bright-field microscopy. Inset: distribution of droplet diameter analyzed after production and after polymerization for a fixed total monomer concentration (*C_T_* = 7.9%). (B) Distribution of droplet diameter for three different batches with three different monomer concentration, *C_T_* = 7.9% (red box), *C_T_* = 9.9% (blue box), *C_T_* = 11.8% (green box).

**Figure S3.**
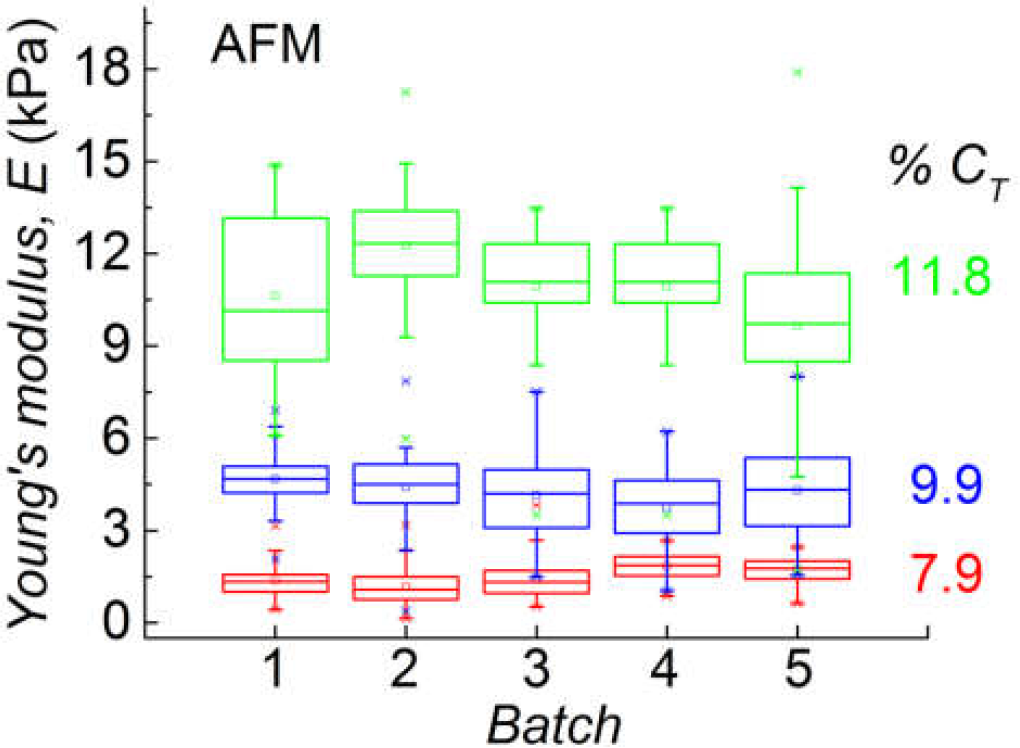
Repeatability of production process and AFM elasticity measurement. (A) Young’s moduli measured by AFM on three groups of five batches. Each group has a different monomer concentration, *C_T_* = 7.9% (red box), *C_T_* = 9.9% (blue box), *C_T_* = 11.8% (green box).

**Figure S4.**
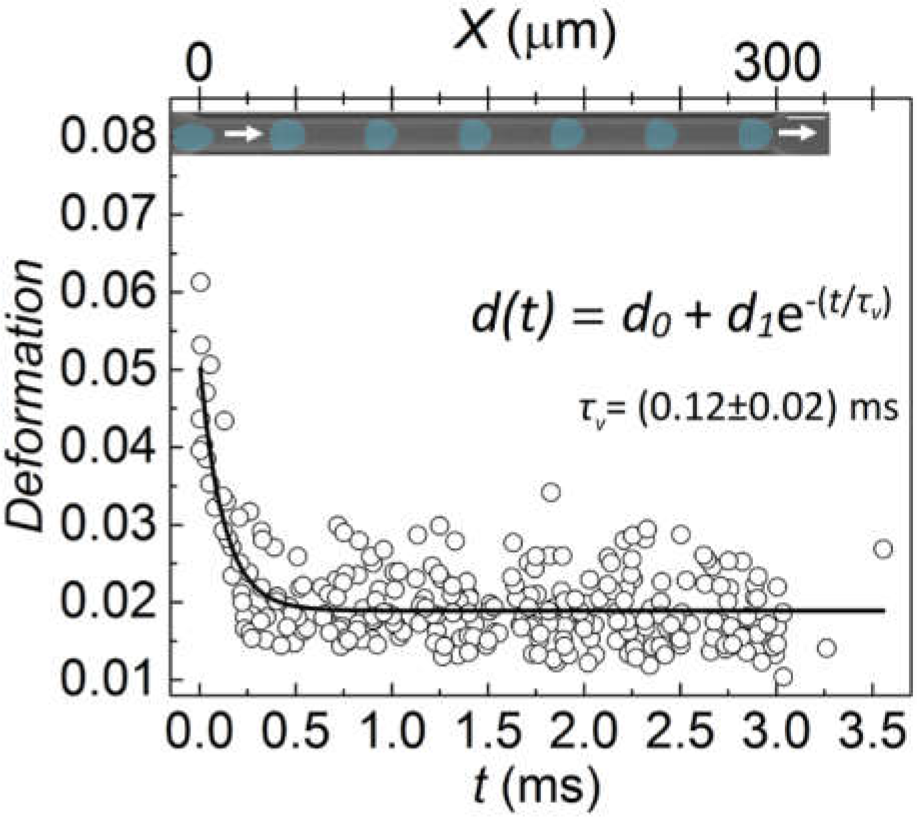
Bead shape evolution in RT-DC microchannel. Bead deformation (*C_T_* = 7.9%) inside the channel depicted over time and the best exponential decay fitting curve (solid line) with *τ_v_* = (0.12 ± 0.02) ms. The inset shows the shape evolution of a bead in the narrow channel (blue mask was applied for better visualization). It was obtained by the superposition of 7 different consecutive frames extrapolated from a video acquired at a frame rate of 2000 fps. Channel length: 300 µm, scale bar: 20 µm.

**Figure S5.**
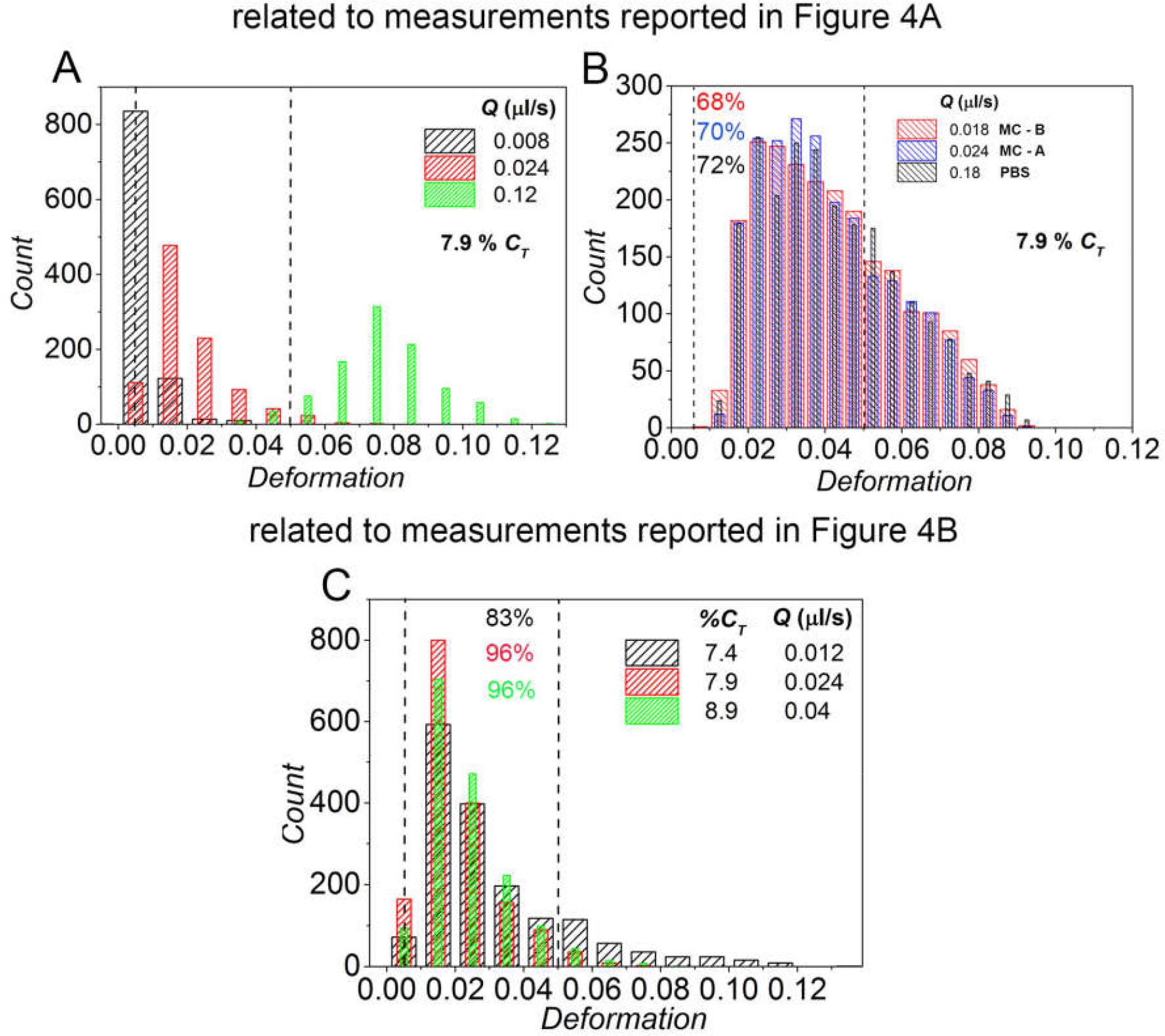
Flow rate optimization for RT-DC measurement, based on bead composition and effective buffer viscosity. The deformation range (0.005 - 0.05) valid for the data analysis is showed with dashed lines. (A) Histogram of PAAm beads (*C_T_* = 7.9%) for three different flow rates and relative deformations: 0.008 µl/s (black bars) with low deformations, 0.024 µl/s (red bars) as suitable settings and 0.12 µl/s (green bars) with high deformations. (B) Histogram of PAAm beads (*C_T_* = 7.9%) using three different measurement buffers: MC-B at 0.018 µl/s (red bars) and 68% accordance with the recommended range, MC-A at 0.024 µl/s (blue bars) and 70% accordance with the recommended range and PBS at 0.18 µl/s (black bars) and 72% accordance with the recommended range. (C) Histogram of PAAm beads for three different polymer concentrations: *C_T_* = 7.4%, flow rate 0.012 µl/s (black bars); *C_T_* = 7.9%, flow rate 0.024 µl/s (red bars); *C_T_* = 8.9%, flow rate 0.04 µl/s (green bars), respectively with 83%, 96% and 96% accordance with the recommended range.

**Figure S6.**
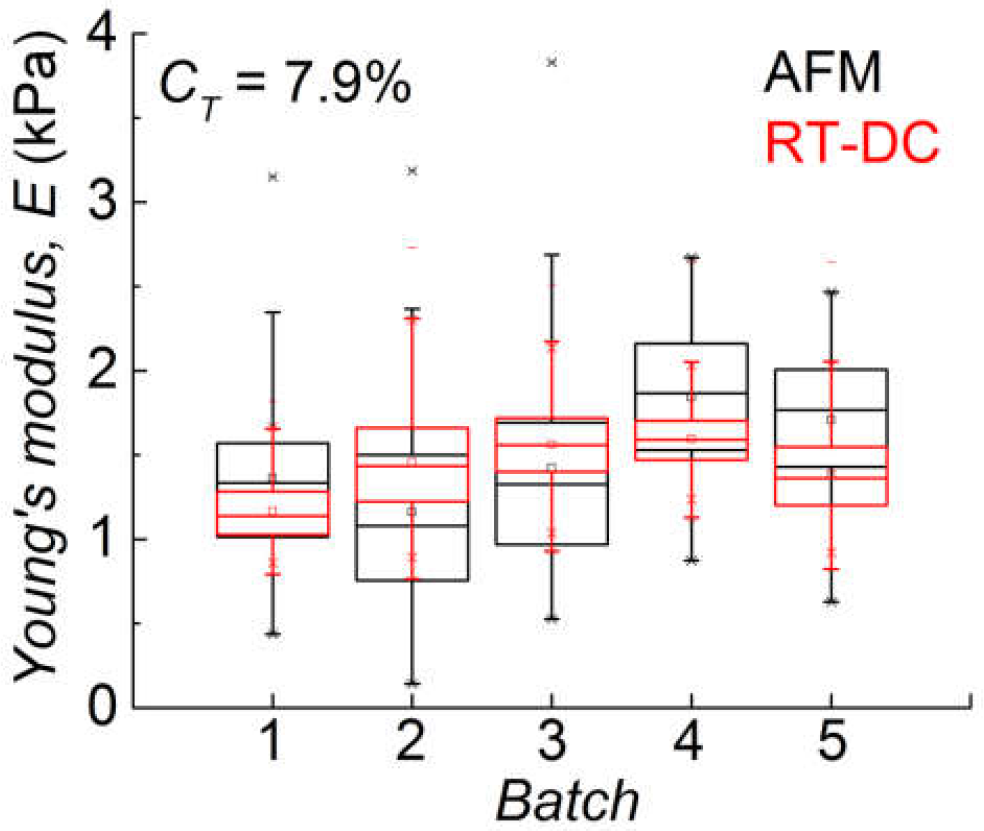
Comparison between AFM and RT-DC measurements. Youn’s modulus obtained with AFM indentation (black boxes) and RT-DC (red boxes) for five different batches produced by using the same monomer concentration (*C_T_* = 7.9%).

**Figure S7.**
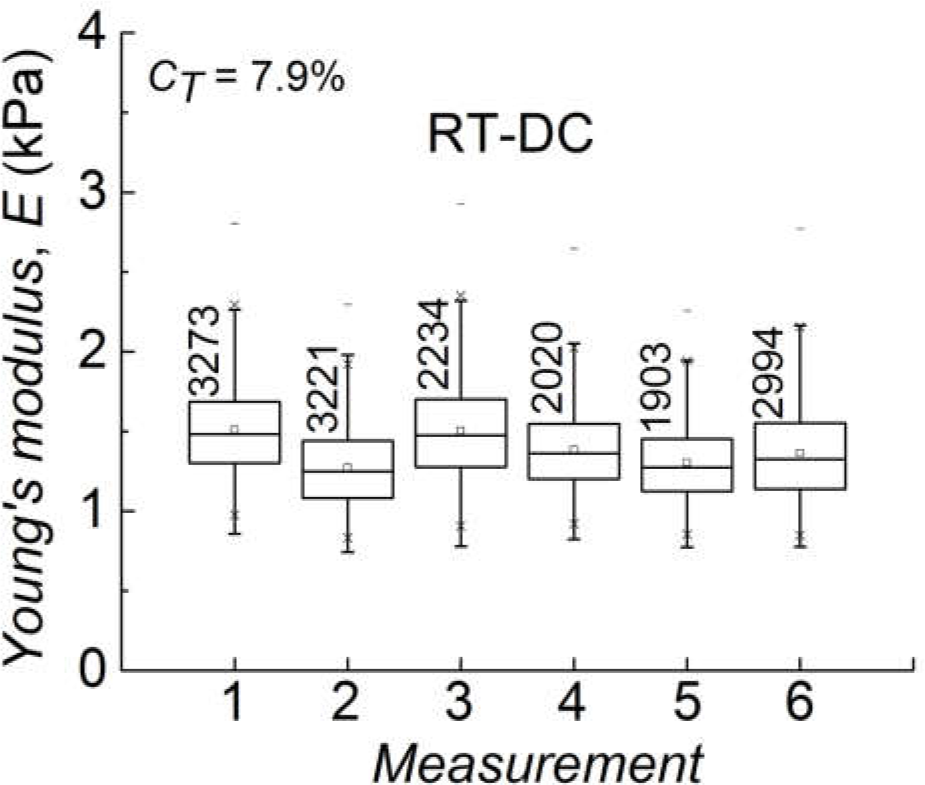
Consistency of RT-DC measurements. Young’s modulus measured by RT-DC six times, during two different days, on the same batch, having a total monomer concentration *C_T_* = 7.9%.

**Table S1.** Diameter mean value for droplets (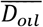) and beads (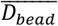), standard deviation *S.D.*, coefficient of variation *C.V.*, for droplets and beads with different monomer concentration (*C_T_*), and the quantification of the bead swelling behavior through the calculation of the normalized volume of equilibrium of the swollen gel *V_eq_* and its standard error (*Δ* (*V_eq_*)). *n* is the number of measured beads.

| Size analysis |  |  |  |  |  |  |  |  |  |  |  |
| --- | --- | --- | --- | --- | --- | --- | --- | --- | --- | --- | --- |
| Droplets |  |  |  |  | Beads |  |  |  |  | Swelling |  |
| $C_T$<br>(%) | $n$ | $\overline{D_{oil}}$<br>($\mu\text{m}$) | S.D.<br>($\mu\text{m}$) | C.V.<br>(%) | Batch | $n$ | $\overline{D_{bead}}$<br>($\mu\text{m}$) | S.D.<br>($\mu\text{m}$) | C.V.<br>(%) | $V_{eq}$ | $\Delta(V_{eq})$ |
| 5.9 | 2789 | 12 | 1 | 8.3 |  | 696 | 18 | 1 | 5.5 | 3.4 | 0.6 |
| 7.4 | 3618 | 11.6 | 0.9 | 7.7 |  | 2596 | 15.7 | 0.8 | 5.1 | 2.5 | 0.5 |
| 7.9 | 2266 | 11.8 | 0.8 | 6.8 | 1 | 1917 | 15.3 | 0.8 | 5.2 | 2.2 | 0.5 |
|  |  |  |  |  | 2 | 1930 | 15.1 | 0.8 | 5.3 | 2.1 | 0.4 |
|  |  |  |  |  | 3 | 521 | 15.2 | 0.5 | 3.3 | 1.8 | 0.4 |
| 8.9 | 2822 | 12.4 | 0.7 | 5.6 |  | 1790 | 15.0 | 0.6 | 4 | 1.7 | 0.4 |
| 9.3 | 3228 | 12.2 | 0.9 | 7.4 |  | 3469 | 14.6 | 0.4 | 2.7 | 1.7 | 0.4 |
| 9.9 | 2716 | 12.5 | 0.9 | 7.2 | 1 | 1914 | 14.3 | 0.6 | 4.2 | 1.5 | 0.4 |
|  |  |  |  |  | 2 | 865 | 14.1 | 0.6 | 4.2 | 1.4 | 0.4 |
|  |  |  |  |  | 3 | 992 | 13.9 | 0.5 | 3.6 | 1.4 | 0.4 |
| 11.8 | 2194 | 12.6 | 0.9 | 7.1 | 1 | 2759 | 12.9 | 0.5 | 3.9 | 1.1 | 0.4 |
|  |  |  |  |  | 2 | 992 | 12.8 | 0.5 | 3.9 | 1.0 | 0.3 |
|  |  |  |  |  | 3 | 811 | 12.6 | 0.5 | 4 | 1 | 0.3 |

**Table S2.**
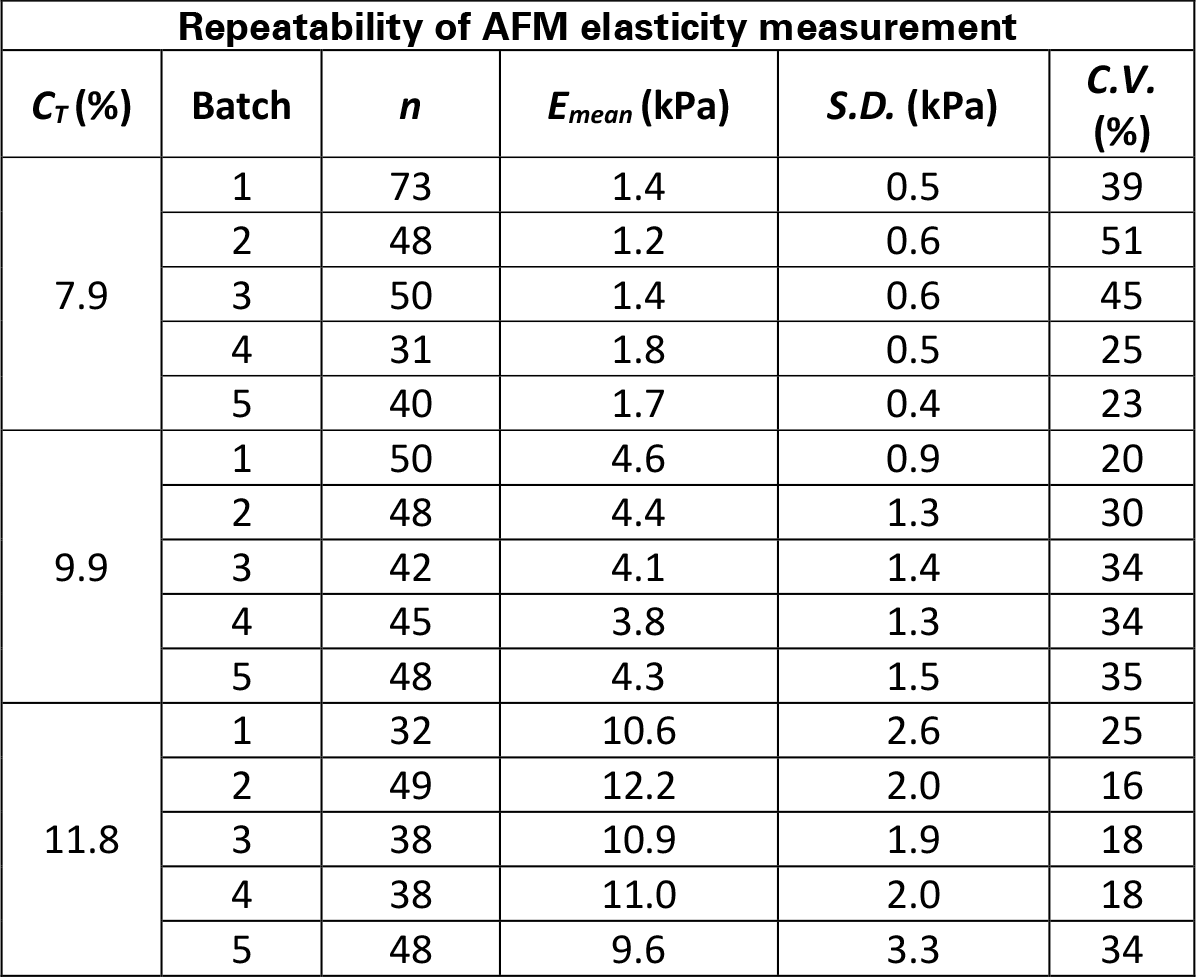
Young’s Modulus mean value *E_mean_*, standard deviation *S.D.*, and coefficient of variation *C.V.*, measured by AFM on three groups of five batches. Each group has a different monomer concentration: *C_T_* = 7.9%, *C_T_* = 9.9%, *C_T_* = 11.8%. *n* is the number of measured beads.

| Repeatability of AFM elasticity measurement |  |  |  |  |  |
| --- | --- | --- | --- | --- | --- |
| $C_T(\%)$ | Batch | $n$ | $E_{mean}$ (kPa) | $S.D.$ (kPa) | $C.V.$ (%) |
| 7.9 | 1 | 73 | 1.4 | 0.5 | 39 |
|  | 2 | 48 | 1.2 | 0.6 | 51 |
|  | 3 | 50 | 1.4 | 0.6 | 45 |
|  | 4 | 31 | 1.8 | 0.5 | 25 |
|  | 5 | 40 | 1.7 | 0.4 | 23 |
| 9.9 | 1 | 50 | 4.6 | 0.9 | 20 |
|  | 2 | 48 | 4.4 | 1.3 | 30 |
|  | 3 | 42 | 4.1 | 1.4 | 34 |
|  | 4 | 45 | 3.8 | 1.3 | 34 |
|  | 5 | 48 | 4.3 | 1.5 | 35 |
| 11.8 | 1 | 32 | 10.6 | 2.6 | 25 |
|  | 2 | 49 | 12.2 | 2.0 | 16 |
|  | 3 | 38 | 10.9 | 1.9 | 18 |
|  | 4 | 38 | 11.0 | 2.0 | 18 |
|  | 5 | 48 | 9.6 | 3.3 | 34 |

**Table S3.** Young’s Modulus mean value *E_mean_*, standard deviation *S.D.*, and coefficient of variation *C.V.*, measured by AFM and RT-DC on five batches produced with the same monomer concentration (*C_T_* = 7.9%). *n* is the number of measured beads.

| Different batches 7.9% $C_T$ | | | | | | | | |
| --- | --- | --- | --- | --- | --- | --- | --- | --- |
| AFM |  |  |  |  | RT-DC |  |  |  |
| Batch | $n$ | $E_{mean}$ (kPa) | $S.D.$ (kPa) | $C.V.$ (%) | $n$ | $E_{mean}$ (kPa) | $S.D.$ (kPa) | $C.V.$ (%) |
| 1 | 73 | 1.4 | 0.5 | 39 | 1607 | 1.2 | 0.2 | 15 |
| 2 | 48 | 1.2 | 0.6 | 51 | 1600 | 1.4 | 0.3 | 21 |
| 3 | 50 | 1.4 | 0.6 | 45 | 1970 | 1.5 | 0.2 | 15 |
| 4 | 31 | 1.8 | 0.5 | 25 | 6534 | 1.6 | 0.2 | 11 |
| 5 | 40 | 1.7 | 0.4 | 23 | 2020 | 1.4 | 0.3 | 18 |

**Table S4.**
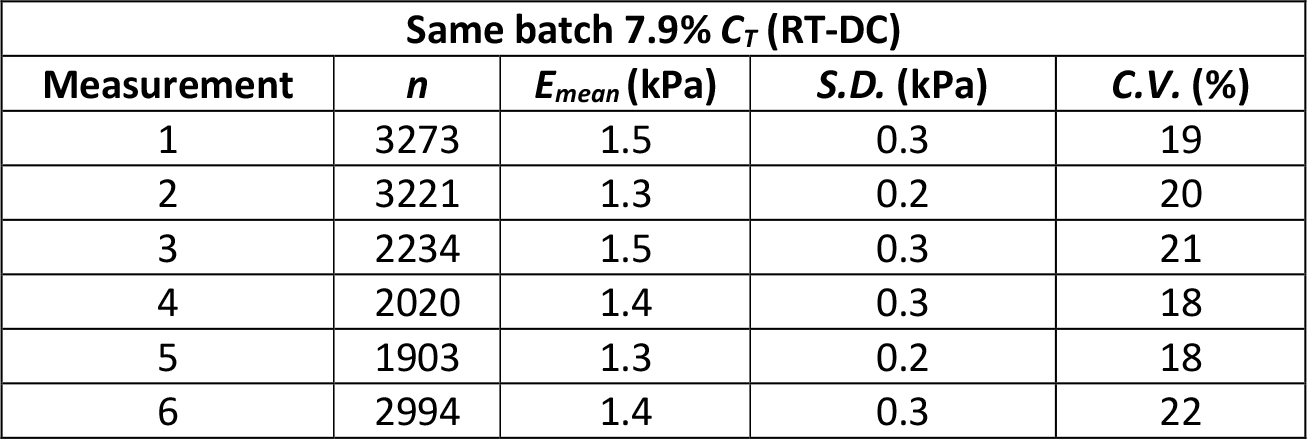
Young’s Modulus mean value *E_mean_*, standard deviation *S.D.*, and coefficient of variation *C.V.*, measured by RT-DC on the same batch, *C_T_* = 7.9%, six times during two different days. *n* is the number of measured beads.

**Table S5.** Young’s Modulus mean value *E_mean_*, standard deviation *S.D.*, and coefficient of variation *C.V.*, measured by AFM on unsorted and sorted microgel beads and by RT-DC on unsorted beads. *n* is the number of measured beads.

| Sorted beads |  |  |  |  |  |
| --- | --- | --- | --- | --- | --- |
| Technique | Sample | $n$ | $E_{mean}$ (kPa) | $S.D.$ (kPa) | $C.V.$ (%) |
| AFM | Unsorted | 50 | 1.2 | 0.5 | 41 |
|  | FSC low | 22 | 0.5 | 0.2 | 37 |
|  | FSC mid | 26 | 0.9 | 0.2 | 23 |
|  | FSC high | 26 | 1.3 | 0.3 | 26 |
| RT-DC | Unsorted | 1402 | 1.1 | 0.2 | 19 |

Video S1 Pre-gel droplet production (*P_oil_* =850 mbar, *PPAAm* =700 mbar). The video was recorded at 3000 fps and showed at 30 fps.

